# Local increases in admixture with hunter-gatherers followed the initial expansion of Neolithic farmers across continental Europe

**DOI:** 10.1101/2024.06.10.598301

**Authors:** Alexandros Tsoupas, Carlos S. Reyna-Blanco, Claudio S. Quilodrán, Jens Blöcher, Maxime Brami, Daniel Wegmann, Joachim Burger, Mathias Currat

## Abstract

The replacement of hunter-gatherer lifestyles by agriculture represents a pivotal change in human history. The initial stage of this Neolithic transition in Europe was instigated by the migration of farmers from Anatolia and the Aegean basin. In this study, we modeled the expansion of Neolithic farmers into Central Europe from Anatolia, along the Continental route of dispersal. We employed spatially explicit simulations of palaeogenomic diversity and high-quality palaeogenomic data from 67 prehistoric individuals to assess how population dynamics between indigenous European hunter-gatherers and incoming farmers varied across space and time. Our results demonstrate that admixture between the two groups increased locally over time at each stage of the Neolithic expansion along the Continental route. We estimate that the effective population size of farmers was about five times that of the hunter-gatherers. Additionally, we infer that sporadic long distance migrations of early farmers contributed to their rapid dispersal, while competitive interactions with hunter-gatherers were limited.

**Teaser:** The first farmers of continental Europe increasingly admixed over time with indigenous hunter-gatherers.

## Introduction

Palaeogenomic studies have revealed that the genetic trajectory of modern Europeans has been strongly shaped by the expansion into Europe of early farmers from the Aegean Basin and western Anatolia starting around 8,600 YBP (*1–3*), introducing agriculture in regions that were previously dominated by hunter-gatherer lifestyles (*4*). The question of whether the Neolithic transition was primarily a cultural process, in which hunter-gatherer groups adopted farming from neighboring communities (*5*), or instead a demic process, in which farmers migrated from Southwest Asia into Europe (*6*), has long been discussed, with studies highlighting the regional role of both processes (e.g. *7*, *8*). This long-standing debate decisively shifted towards favoring migration as a key factor following Bramanti *et al*.’s work (*1*), which revealed substantial genetic differences between Central European hunter-gatherers and the first agriculturalists. Indeed, the latter displayed a genetic signature that only emerged in Europe during the Neolithic period. Subsequent genomic studies supported these findings (e.g. *9*), successfully tracing the ancestral roots of early Neolithic populations in Europe to the area encompassing Western Anatolia and the Aegean basin (*2*, *3*). It was further established that the Early Neolithic inhabitants of present-day Iran differed genetically from Aegean and European farmers, demonstrating that the Neolithic migration chain did not begin in the Fertile Crescent as some researchers have long assumed (*10*). Recently, Marchi *et al*. presented evidence that the European early Neolithic population originated in Anatolia through a process involving population mixing during the Last Glacial Maximum and genetic drift at the onset of the Holocene. The resulting population adopted agriculture in Anatolia and gradually expanded towards Central Europe, eventually reaching Northern and Western Europe (*11*). This dispersal route, proceeding along the Danube, the Rhine and other major river valleys, is described in the archaeological literature as the “Continental” or “Danubian route” of Neolithic expansion (e.g. *7*, *12*). It is usually contrasted with the “Maritime” or “Mediterranean” route. The latter followed a coastal path along the Mediterranean shorelines, resulting in the colonization of southern Europe and eventually the settlement of southern France, northern Africa and the Iberian Peninsula (*13*).

The present study focuses on the Continental route of Neolithic expansion. Before the disappearance of hunter-gatherer lifeways in Europe, farmers and foragers coexisted for many generations and in certain instances for thousands of years (*14*). Many questions remain about the biological interactions between the two populations, which spanned more than 3,000 years, from the Balkans to Denmark (*15*). Although the general pattern of early farmer migration seems consistent across the Western part of Europe, there is regional and temporal variation in the levels of admixture with hunter-gatherers (e.g. *16*–*18*), and regional studies provide valuable insights into local population interactions (e.g. *19*–*21*). Some palaeogenomic data support the assertion that hunter-gatherers and farmers only admixed sporadically at the outset of the Neolithic (*3*, *22*). Other studies indicate that admixture intensified at later stages of the Neolithic (starting approximately 7,000 YBP, *17*, *18*, *21*, *23*), possibly after the so-called ‘crisis’ of the LBK (Linearbandkeramik). This period, which spans until the Central European Middle Neolithic, is characterized by massacre sites, mass graves and widespread abandonment of sites (*24*, *25*). Spatially, the levels of hunter-gatherer ancestry in Neolithic populations appear to increase east of the axis that goes from the Black Sea to the Baltic region (*15*). There is no such clear trend from Southern to Northern Europe, although more admixture at higher latitude was proposed (*18*), possibly linked to a slowdown of the Neolithic expansion speed (*26*). Moreover, Rasteiro & Chikhi (*27*) have described a trend of decreasing female and male genetic lineages of Neolithic farmers in the genome of modern Europeans, with distance from the Middle East, an observation compatible with the demic diffusion model (*6*) including only very limited local hunter-gatherers introgression into migrating farmers at each step of their progression (*28*, *29*).

In the present study, we wanted to better understand the population dynamics underlying the interactions between hunter-gatherers and farmers along the Continental route of the European Neolithic. We modeled the Neolithic expansion by jointly simulating demography, migration, and biological interactions (in the form of admixture and competition) between two populations, within a spatially explicit framework. We then used an Approximate Bayesian Computations (ABC) framework to infer key demographic parameters by comparing the resulting simulated palaeogenomic data to recently published high quality genomic data from 67 individuals. Compared to previous demogenomic modeling of the Neolithic using ancient DNA, our simulation framework is able to manage more explicit spatial features than Marchi *et al.* (*11*) and to handle the spatio-temporal variation of admixture between hunter-gatherers and farmers with more flexibility than Rasteiro & Chikhi (*27*) and Silva *et al.* (*30*). The paleogenomic dataset we used is particularly well suited for demogenomic inferences as it is composed of genome-wide sequences selected from presumably neutral regions (*31*). Moreover, their high sequencing quality allowed us to directly compute molecular indices of intergenomic diversity (i.e. nucleotide diversity) rather than summarizing the molecular information through estimated admixture proportions from putative source populations (e.g. *32*). We specifically examined what dynamics characterized the demographic processes underlying the interactions between hunter-gatherers and farmers and whether variations in admixture between these populations are detectable over time and space.

## Results

### Model choice among five demographic scenarios

We started by comparing 5 variations of the model (thereafter called scenarios) in which admixture with indigenous hunter-gatherers (HG) varied through space and/or time when Neolithic farmers (FA) migrated from Anatolia towards Europe. The assimilation rate in the investigated scenarios was constant or increasing in time, while it was constant, increasing, or decreasing with distance from Anatolia (see Fig. 1D for a graphical description of the 5 scenarios). It is worth noting that in all scenarios, only a relatively small proportion of simulations (∼4%) were able to replicate the entire dataset of samples. The number of simulations available for the ABC estimation is thus relatively low despite intensive computation. We therefore used the random forest ABC for model choice, as Supplementary Text 1 demonstrates that using only 13,000 simulations per scenario is enough to differentiate them to an extent when summarizing the genomic information into pseudo-haploid sequences. All five investigated scenarios were able to reproduce the observed molecular diversity (Table S1), as shown by marginal density p-values larger than 0.05 and were therefore retained for further analysis.

**Fig. 1.**
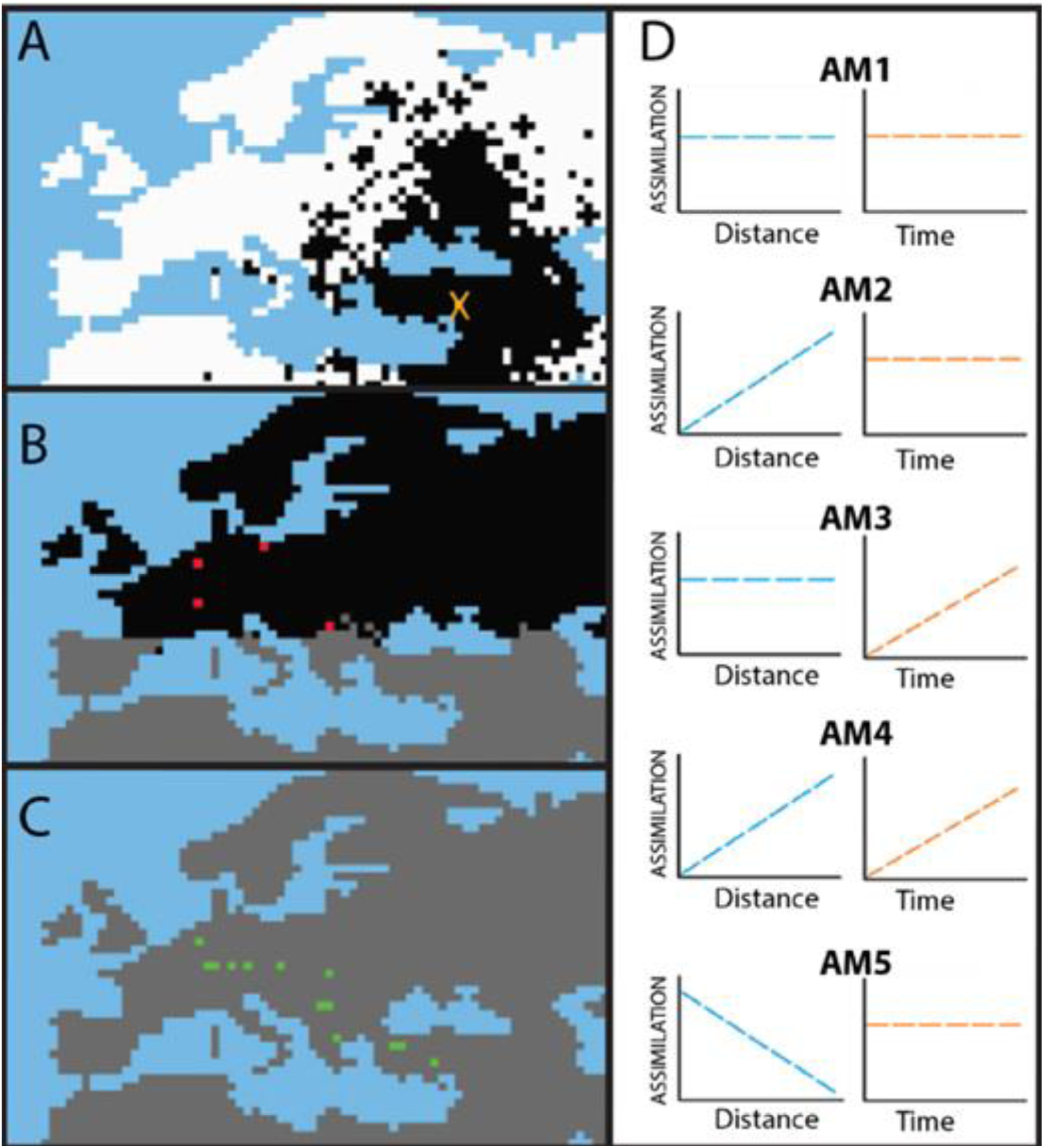
Graphical representation of spatially explicit simulations and investigated scenarios. Blue = water cells; white = cells with only HG; black = cells with both HG and FA; gray = cells with only FA. **(A)** Example of simulation with the spread (including Long Distance Dispersals) of FA from the orange cross towards Northwestern Europe through the Continental route. **(B)** Two zones with different competition coefficients. In the gray zone, α_S_ = 1, while in the black zone, α_N_ is variable (taken from a prior distribution) allowing a longer cohabitation between HGs and FAs; red cells show demes where HG samples are simulated. **(C)** At the end of the simulation, HGs disappear from the whole simulated area; green cells show demes where FA samples are simulated. **(D)** Graphical representation of the differences of the investigated scenarios. The assimilation rate (*γ)* of HGs into FAs can be constant or increase with time, and be constant, increase, or decrease with distance from the starting point of the FA expansion in Anatolia.

The most probable scenario among the five was AM3 (Table S2), with a posterior probability of 43%. This suggests that the observed levels of genetic differentiation between HGs and FAs are more accurately described by a scenario in which the rate of admixture between the two groups increases locally over time, while its maximum rate remains constant along the expansion axis. However, the probability of misidentifying it as AM4, another scenario with increasing admixture over time is substantial (34.3%, Fig. 2).

**Fig. 2.**
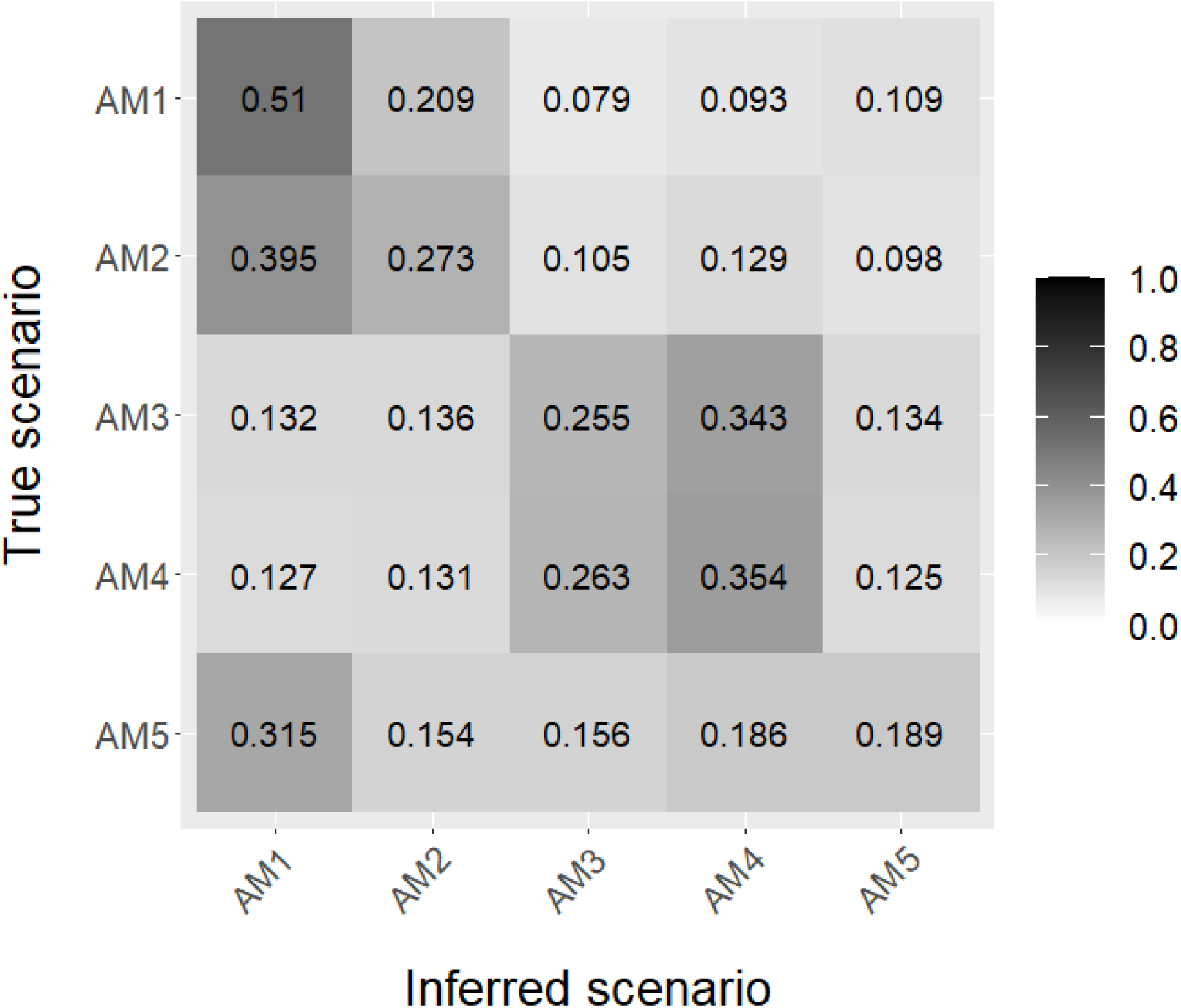
Confusion matrix for the five investigated scenarios. Calculation performed with abcrf R package (random forest approach, *76*), using 33,500 simulations per scenario. The numbers correspond to the proportion of simulations that was assigned to each case by ABC. For the analysis, the untransformed pairwise values of inter- and intra-sample pseudo-haploid nucleotide diversity were used.

Given that individual assigning to the HG or FA groups can affect the results, we chose to rerun simulations removing genomes from the exceptional site of Lepenski Vir in Serbia, in which the group attribution is especially challenging (*19*, *20*). However, it does not change model choice, with model AM3 still being the best with a relative probability of 40% (Supplementary Text 2).

From the confusion matrix (Fig. 2), we observe that scenarios with spatially increasing admixture (i.e. higher admixture rate in the northwest of the Continental route than in the southeast) cannot be differentiated well from the ones where admixture is constant in space (AM2 and AM4 from AM1 and AM3, respectively). However, scenarios in which admixture increases temporally in each deme since their first colonization are easier to differentiate from those in which admixture is constant over time (AM3 and AM4 from AM1 and AM2). Scenario AM5, in which admixture is lower in the Southeast than in the Northwest of the Continental route, produces a pattern of diversity not distinguishable from the other scenarios, as its simulations are randomly classified among the five scenarios. Seeing that scenario AM5 is misclassified more often than correctly identified and that scenarios AM3 and AM4 are hardly distinguishable, we performed a second model choice using a subset of only the three first scenarios (AM1, AM2 and AM3) to avoid noise (Supplementary Text 3). In this case, the most probable scenario is still AM3, with the posterior probability increasing to 52% (Table S7), and a 65.7% probability of being correctly identified (Fig. S6).

### Parameter estimation using a single demographic scenario

The most probable scenario is thus AM3 representing the admixture rate increasing with time in each deme, but in a spatially homogeneous manner. However, our model choice procedure did not definitively rule out the other scenarios (Fig. 2). Therefore, we estimated the various parameters using an additional scenario, SM6, which encompasses the characteristics of all five scenarios with a nuance: in SM6 the duration of the increase of the admixture rate varies within the cohabitation period between HGs and FAs (in contrast to AM3’s constant increase over the whole cohabitation period). This allows us to estimate the duration of the admixture rate increase.

SM6 had a very strong demographic filter, with about 95% of the simulations being rejected because they were not able to reproduce the whole dataset’s characteristics. The marginal density p-value for the remaining simulations under SM6 (p=0.29 with δ = 0.01,0.32 and 0.18 for δ = 0.005 and 0.05, respectively), indicates its capability to reproduce the observed values of molecular diversity. The characteristics of the posterior distributions of the estimated parameters are presented in Table 1 (with δ = 0.01, while results for δ = 0.05 and 0.005 are presented in Table S3), while a graphical representation of the posterior and prior distributions of each parameter is given in Fig. 3. For all parameters except the sequencing error rate, we plotted in addition the distribution of parameter values of the simulations retained by the demographic filter (see green lines in Fig. 3). We call it “demographic posterior”, to differentiate it from the posterior that results from the ABC estimation made on molecular statistics, which we call here “molecular posterior” (see red lines in Fig. 3). For the two *γ* variables (*γ_S_* and *γ_N_)* and the two *K* variables (*K_HG_* and *K_FA_*), we estimated two-dimensional posteriors and plotted the estimated HDIs (Fig. 3), in order to investigate if there are combinations of these variables that are more probable than others.

**Table 1.** Characteristics of the posterior distributions of the estimated parameters of scenario SM6. We performed 100000 simulations with 1% tolerance level, retaining 1000 simulations. The estimation was performed with ABCtoolbox2 (*77*).

| Parameter | Posterior Mode | Posterior Mode relative bias | Posterior Mean | Posterior Mean relative bias | Posterior Lower HDI95 | Posterior Upper HDI95 |
| --- | --- | --- | --- | --- | --- | --- |
| Admixture rate in Southeast Continental route ($\gamma_S$) | 0.024 | 3.96 | 0.049 | 3.67 | 0.003 | 0.096 |
| Admixture rate in Northwest Continental route ($\gamma_N$) | 0.029 | 4.38 | 0.046 | 3.77 | 0.002 | 0.093 |
| Number of generations of $\gamma$ increase ($t_{inc}$) | 70 | 1.49 | 72 | 2.29 | 2 | 136 |
| HG effective population size ($K_{HG}$) | 343 | 0.14 | 357 | 0.12 | 273 | 435 |
| FA effective population size ($K_{FA}$) | 1479 | 0.18 | 1769 | 0.17 | 1230 | 2322 |
| Migration rate of FAs ($m_{FA}$) | 0.4232 | 0.40 | 0.344 | 0.37 | 0.162 | 0.5 |
| Proportion of Long-Distance Dispersals ($P_{LDD}$) | 0.0192 | 2.26 | 0.013 | 3.50 | 0.002 | 0.024 |
| Competition coefficient in Northwest Continental route ($\alpha_N$) | 0.2033 | 0.17 | 0.231 | 0.19 | 0.15 | 0.327 |
| Sequencing Error rate ($\epsilon$) | 0.00006 | 0.22 | 0.00007 | 0.22 | 0.00005 | 0.0001 |

**Fig. 3.**
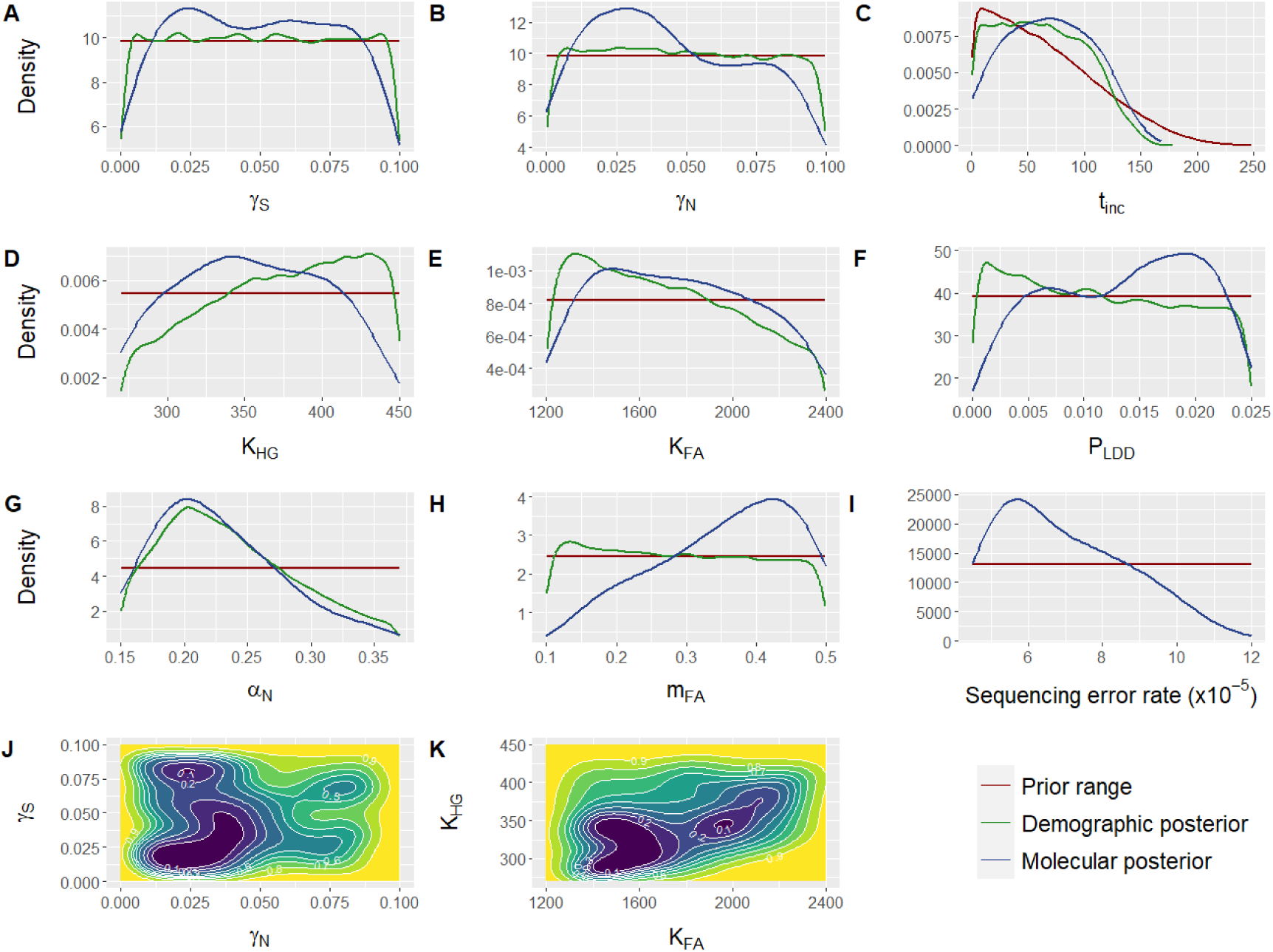
Distribution of the estimated parameter values. In panels A-I, the red line corresponds to the prior distribution, the green one to the distribution of parameter values of the simulations that were able to reproduce the observed dataset (demographic posterior), and the blue line corresponds to the posterior resulting from the ABC estimation (molecular posterior). **(A)** Assimilation rate along the Southeast Continental route (*γ_S_)*; **(B)** Assimilation rate along the Northwest Continental route (*γ_N_)*; **(C)** Number of generations during which the assimilation rate increases (*t_inc_*); **(D)** Effective population size of HGs (*K_HG_*); **(E)** Effective population size of FAs (*K_FA_*); **(F)** Proportion of FA migrations that are Long Distance Dispersals (*P_LDD_)*; **(G)** Competition coefficient along the Northwest Continental route (*α_N_*); **(H)** Migration rate of FAs (*m_FA_*); **(I)** Sequencing error rate (ε); **(J)** Two-dimensional posterior distribution of assimilation rate (*γ*) along the Southeast Continental route (*γ_S_)* against the assimilation rate along the Northwest Continental route (*γ_N_)*; **(K)** Two-dimensional posterior distribution of effective population size (*K*) of HGs (*K_HG_*) against the Effective population size of FAs (*K_FA_*). For the two-dimensional posteriors, yellow represents the value combinations with the lowest probability and blue the combinations with the highest one.

In line with the model selection process, scenario SM6 does not suggest any spatial variability in the admixture rate. *γ* in both parts of the Continental route is characterized by wide, overlapping posterior ranges (*γ_S_*=[0.3%-9.6%] and *γ_N_* = [0.2%-9.3%]), with posterior modes at 2.4% and 2.9% respectively (Table 1). The mean and mode are characterized by large relative biases (Table 1), which implies that these point estimates are not accurate. The two-dimensional posterior did not indicate any particular combination of *γ_S_* and *γ_N_* being more probable than others (Fig. 3J). From the two-dimensional posterior we estimated that γ*_S_* being larger than γ*_N_* and the reverse are equally probable (48.3% and 51.7%, respectively). Moreover, we estimated that the admixture rate from HGs to FAs was increasing for about 70 generations on average (∼1,750 years), with the values varying between 2 and 136 generations (∼50-3,400 years), the point estimate being relatively imprecise (Table 1).

The effective population sizes (*K*, in number of chromosome copies) have broad posterior ranges compared to the prior ones (Table 1). However, the mean and mode relative biases are low, showing relatively accurate means of 357 gene copies for HGs and 1,769 for FAs. Also, the posterior distribution of *Kfa* (Fig. 3E) is not uniform but skewed towards lower values. Based on the two-dimensional posterior (Fig. 3K), we estimated that the ratio of the effective population sizes (*Kfa /Khg*) ranges between three and seven times larger in favor of FAs, with a probability of 96%, and a mean of five times.

Our findings indicate that the presence of a low level (∼2%) of Long Distance Dispersals (LDD) provides a better explanation of the data as 0 is excluded from the posterior range [0.2-2.4%]. This suggests that sporadic LDD events play a role in managing the rapid Neolithic expansion. When using shorter LDD events, of 300 km on average, the marginal density p-value of the simulated statistics is lower than 0.05, which means that the scenario is no longer able to reproduce the observed summary statistics. Additionally, under a scenario without LDDs, only 0.03% of the simulations passed the demographic filter. Altogether, these observations suggest that sporadic LDD events are needed to cope with the fast Neolithic expansion and for developing the observed pattern of genetic diversity measured in our study. Nevertheless, the impact of their frequency on the measured genetic diversity in our study appears to be limited as the point estimate lacks precision (wide posterior range relative to the prior distribution and the large relative bias).

We calculated the competition coefficient *α_N_* to range approximately from 15% to 33%, a narrower range compared to the original 0-100% range (Supplementary Text 4). Both the mode and mean values fall around 20-23%, demonstrating low relative bias (Table 1), indicating accurate estimation of this parameter. The posterior distribution of the sequencing error rate points to a mode value of 5.7e-05 with low relative bias, corresponding to an NGS quality score of 42.

Even after excluding Lepenski Vir genomes from the analysis, for which the group attribution is challenging, the parameter estimation remains largely unchanged, confirming the robustness of our results (Supplementary Text 2).

## Discussion

### Local temporal increase in the admixture between HGs and FAs

The demographic modeling approach presented here underscores a nuanced demic diffusion model for the Neolithic spread from Northwest Anatolia extending through the Balkans into Central Europe and yields novel demographic parameter estimates. These findings corroborate a temporal dynamic in admixture between HGs and FAs, indicating a gradual rise in genetic exchange within each location (e.g., deme) over successive generations of cohabitation between these groups. Our results thus suggest that the resurgence of HG ancestry in Neolithic populations after ∼7,000 YBP (*17*, *18*, *21*) is best explained by local increases in admixture between HGs and FAs over time, rather than a constant accumulation of HG ancestry. It should be noted that our model does not test admixture pulses (*16*), defined as two periods of admixture separated by a period of genetic isolation (*33*), which could be a future development of our approach.

It is important to emphasize that the estimated admixture rate is not equivalent to the proportion of HG ancestry in Neolithic populations. Equating the observed proportion of HG ancestry in farming populations with the frequency of admixture can result in misconceptions about historical interactions, as demonstrated by simulation studies (*29*). Instead, the inferred admixture rate offers a more precise measure of the actual interactions between these groups. For instance, theoretical simulations have shown that a low admixture rate (e.g. 5%) at each stage of the Neolithic expansion can lead to a large HG ancestry (e.g. 50%) due to the cumulative effect of population migration, demography and biological interactions (*29*).

Our estimates show that admixture increased over time, but its magnitude remained low. This ranges from no gene flow upon arrival of farming communities in a local population (deme), to approximately 5% (with a maximum of 10%) of interactions between hunter-gatherers (HG) and farmers (FA) leading to admixture during later phases of the Neolithic. The “intensity” of admixture - the 95 HDI of the rate we estimated (0.2%-9.3%) - is of the same order as the values (2%-7%) of HG incorporation in the first farming communities estimated under a different (not spatially explicit) model by Marchi *et al*. (*11*). Moreover, it falls within the range (< 10%) estimated with spatially explicit simulations by Silva *et al.* (*30*) based on ancient mitochondrial DNA and is compatible with the estimation made from autosomal SNPs by Silva *et al.* (mean of ∼8.8%, 90 HDI = 5-14%, *34*). We refrained from further comparisons to previous studies, given 1) the differences between the datasets analyzed (>320 samples for a single sex-specific locus in *30* and only two whole genomes in *34*); 2) the difference between the admixture model implemented (constant admixture in *34* and *11* but variable in *30* and here); 3) the limited precision of our admixture rate estimation (wide posterior distribution). This relatively low precision may be due in part to the stochastic effect of LDDs, which has been shown to diminish introgression of local (e.g. HG) alleles during (e.g. FA) range expansions (*35*).

Our results suggest that early farming communities quickly settled into favorable environments suited for agriculture, potentially occupying distinct spatial territories from HG groups (*4*, *36*). This spatial and possibly cultural separation likely restricted interactions between these groups. Subsequently, as agricultural practices stabilized, acculturation or genetic exchanges with external HGs became more intensive. The ‘crisis’ in LBK farming communities around 7,000 BP, which resulted in a significant population reduction (*37*, *38*), may have contributed to a steady increase in the genetic contribution of hunter-gatherers to local farming communities. The delayed admixture scenario, supported by our results, draws attention to what prehistorian Stuart Piggott once described as “secondary Neolithic” cultures (*39*). Such cultures, initially identified in Western Europe based on continuities in chipped stone traditions, are traditionally thought to represent the assimilation of Neolithic features by hunter-gatherer-fishers after the first impact of the arrival of immigrant farmers in the region (*39*).

Our model does not allow the identification of specific cultures or factors influencing this increased assimilation, whether they are ecological, economic or climate-related (*37*, *40*). Moreover, it remains unclear whether increased interactions between foragers and farmers led to widespread adoption of Neolithic practices, as assumed by traditional frontier expansion models in archaeology, such as Marek Zvelebil’s three-stage “availability” model (*41*). We estimated the increase of admixture to have lasted a long time period (∼1750 years), consistent with the prolonged coexistence estimated in some parts of Central and Northern Europe (*14*, *42*). Our estimation tends however to be longer than the admixture time of 500-840 years estimated for Hungary and 280-700 years for Germany (*18*).

### No indication of spatial variation in the admixture between HGs and FAs

Unlike the temporal aspect, our findings do not reveal spatial variability in the admixture rate between HGs and FAs along the Continental Neolithic route. Neither a scenario suggesting a higher admixture rate in the northwestern part of the Continental route (AM2), nor one indicating a lower rate (AM5), corresponds better to the observed data than a scenario with an equal rate (AM1). Moreover, the admixture rates estimated in the Northwest and in the Southeast of the Continental route are not different, with largely overlapping 95HDI (Table 1). Even if it aligns with previous demogenetic modeling studies that relied on a spatially homogeneous admixture rate at the continental scale (*27*, *29*, *43*, *44*), this is in contrast to the hypothesis posited by Betti *et al*. (*26*), which proposes a higher admixture rate in the Northwest.

Our approach has been tailored to identify overarching spatial trends along the Continental route. However, a Neolithic expansion model based on radiocarbon dates, as noted by Fort (*7*), has highlighted regional variations in the demographic or cultural diffusion of the Neolithic. So, we acknowledge the possibility that our analysis may not capture nuanced regional variations. Moreover, our cross-validation analysis revealed that our approach is better suited to identify temporal variation in admixture than spatial variation, for which we may lack power. Distinguishing between equivalent scenarios, with or without spatial increase in admixture (AM1 vs AM2 vs AM5 and AM3 vs AM4 in Fig. 2), remains for instance challenging.

### Demic diffusion with sporadic long distance migrations

Our results support the view that most of the migration movements occurred in a stepping-stone manner, with a very small proportion involving long distance migration (LDD), contributing to the rapid advance of the Neolithic expansion front along the Continental route in the Balkans and Central Europe (e.g. *26*, *45*). This is in accordance with two modeling studies based on radiocarbon dates in the Central Balkans, supporting a rapid demic diffusion model of expansion with sporadic leapfrog migration events (*46*), possibly including some LDD occurring behind the expansion front with groups of migrants traveling between 88 and 150 km before settling down (*47*). In comparison, we set the average LDD distance in our study to about 800 km, in order to contrast their effect with short-scale migrations that occurred 100 km away (the size of a deme). Using a shorter LDD distance of about 300 km is not able to reproduce the molecular diversity in our dataset, having marginal density p-values < 0.05, thus confirming that a low amount of LDD is needed.

The detection of sporadic LDD in our model contrasts with results from Marchi *et al*. (*11*), who did not find a signal for LDD along the Continental route. However, both results are not necessarily incompatible, given the very low proportion of LDD migrations we estimated in our study and the differences in the models used. Indeed, the simulations performed by Marchi *et al*. used a much lower number of demes (<10) compared to ours (>100 between the most distant samples) and it is harder to detect rare long distance events of migration at lower spatial resolution. Altogether, our results support the view that the Neolithic expansion along the Continental route consisted of sporadic long distance dispersals similar to the ‘leapfrog colonization’ model suggested by archaeologists (*8*, *48*), with short scale migrations making up the bulk of the demic diffusion pattern.

### Limited population competition and population effective size ratio

Studies taking into account interpopulation competition between HGs and FAs remain rare. While prior modeling studies integrated competition between HGs and “converted” FAs using the “wave of advance” numerical model (*49*, *50*), as well as competition between HGs and “initial” FAs (*51*, *52*), they primarily focused on how competition affects the speed of the advancing FA wave. None of these studies attempted to quantify competition based on molecular data, as we have done in our research. We estimated that a low competition rate between HGs and FAs is necessary to reproduce the long cohabitation period between them in Central Europe (*14*, *42*), together with a specific ratio of population sizes for the two groups of ∼5 fold. We estimated a coefficient of competition of ∼20% under a Lotka-Volterra model, which could be interpreted as both populations occupying partially overlapping niches, i.e. by exploiting about one fifth of resources in common. Some potential explanations for the relatively low competition could be that the early migrating FAs occupied areas not inhabited by HGs (*36*) or that the arrival of FAs in Central and Northern Europe affected the yield of their crops and therefore their competitiveness (*26*).

In combination with the limited competition rate, we estimated the effective population size of FAs to be between three and seven times higher than that of the HGs. This ratio is smaller than the value (10x-30x) assumed in previous simulation studies of the Neolithic transition (*6*, *29*, *43*, *44*) and within the range used in Silva *et al*. (*30*). By dividing our estimated effective size (in number of chromosome copies) by two to get the number of effective individuals and then multiplying the result by three (*53*), we were able to roughly estimate a FA census size of approximately 0.27 individuals/km^2^ (95 HDI = 0.19-0.35) within a deme of 100×100 km^2^. This estimate is close but lower than the estimate of Silva *et al*. (*30*), based on ancient mitochondrial data (0.46 individuals/km^2^) and in line with the upper range (0.6 individuals/km^2^) estimated by Zimmermann *et al.* (*54*) for the Linear Pottery culture (LBK).

### Model adaptations and limitations

The spatially explicit simulator, SPLATCHE3, had to be adapted to the specificities of the palaeogenomic dataset analyzed. In particular, we incorporated a sequencing error rate in our simulations, as it may bias the inferences when not taken into account (*55*). The posterior distribution points to a sequencing error rate of 5.7e-05, which is lower than the rate usually admitted for NGS: around 1e-03 (*56*) to 2.4e-03 (*57*) and within the range estimated for NGS data after computational correction (1e-05 - 10e-05, *58*). This low estimated error rate was expected, since we calculated pairwise distances between pseudo-haploid genome calls and only at sites covered at least twice after trimming away bases potentially affected by post mortem damage.

Although our modeling framework is a step toward considering non uniform demic models of Neolithic expansion by varying admixture rates between HGs and FAs at different latitudinal stages of the Continental route, there are still some missing features that would be worth investigating in the future. First, while our model uses logistic growth, we assumed that the carrying capacity of HGs and FAs was uniform and constant over both time and space. We thus did not consider the inferred fluctuations in FA population sizes during the Neolithic (*38*, *54*, *59*), nor the differential occupation of diverse environments by both Mesolithic HGs (*60*, *61*) and Neolithic FAs (*48*, *62*). Second, by using uniform and constant migration rates, we did not consider arrhythmic leaps in the Neolithic spread (*63*), as well as potential faster spread along the coastlines and rivers (*64*). We can imagine, however, that such arrhythmias would have a larger impact on spatial molecular patterns than on the temporal pattern we identified in our study. Furthermore, incorporating the estimated age range for each observed genome instead of relying solely on mean values could enhance the accuracy of the demographic filter, particularly concerning parameters associated with population movement. Despite leveraging a high-quality palaeogenomic dataset for demographic inferences, our study faced hurdles in differentiating scenarios and attaining accurate parameter estimates. These challenges stemmed from the potential for similar outcomes arising from different combinations of parameter values (equifinality issue), alongside information loss during data conversion and dimensionality reduction. These complexities highlight the delicate balance and trade-offs inherent in extracting information about past population dynamics from ancient DNA.

Our simulation framework establishes a foundational platform for the future development of more detailed scenarios to study how population demographic variations, migrations, and interactions affected spatiotemporal patterns of genomic diversity. As new palaeogenomic data continue to accumulate with improved sequencing quality, this iterative process can significantly enhance our understanding of ancient population dynamics. This will enable more refined demographic estimates regarding competition and admixture along the Neolithic Continental route, but also other major events that have affected the history of our ancestors at different periods.

## Materials and Methods

### Dataset

To investigate the expansion of Neolithic farmers and their interaction with European hunter-gatherers, we analyzed high coverage genomic data from 67 ancient individuals located along a transect between north-western Anatolia and Central Europe, from a total of 17 archaeological sites (Fig. 4). The majority of the dataset under analysis comprises genomes from Hofmanová *et al*. (*20*), encompassing 49 high-quality target enrichment captures of neutral genomic regions spanning a total of 5 Megabases referred to as “neutralomes” (*31*). Per design they should allow more accurate inferences about demographic events by minimizing bias resulting from functional restrictions or background selection. This dataset was complemented by 18 published genomes of equivalent quality and coverage fitting the geographic and temporal requirements of our study (Supplementary Data S1). The raw data of the neutralomes and genomes were processed as described in Hofmanová *et al.* (*20*). The dataset consists of 16 Paleolithic, Mesolithic and Neolithic hunter-gatherers (HGs) and 51 Neolithic farmers (FAs). Group assignments of each genome as HG or FA were taken from the original publications, based on criteria such as the presence or absence of food-producing activities at the site. While the distinction between early Holocene hunter-gatherers and farmers is relatively clear-cut in continental Europe, the picture is more contrasted to the east. For instance, at 11^th^/10^th^ millennium BP Boncuklu, in Central Anatolia, small-scale farming was practiced next to hunting and gathering (*65*). At 9^th^ millennium BP Lepenski Vir in the Danube’s Iron Gates, incoming farmers appear to have adopted fishing (*19*). The estimated age of the HG genomes ranges between 13,700 and 5,719 YBP, while the age of FAs is between 10,078 and 5,368 YBP (see Supplementary Data S1 for details).

**Fig. 4.**
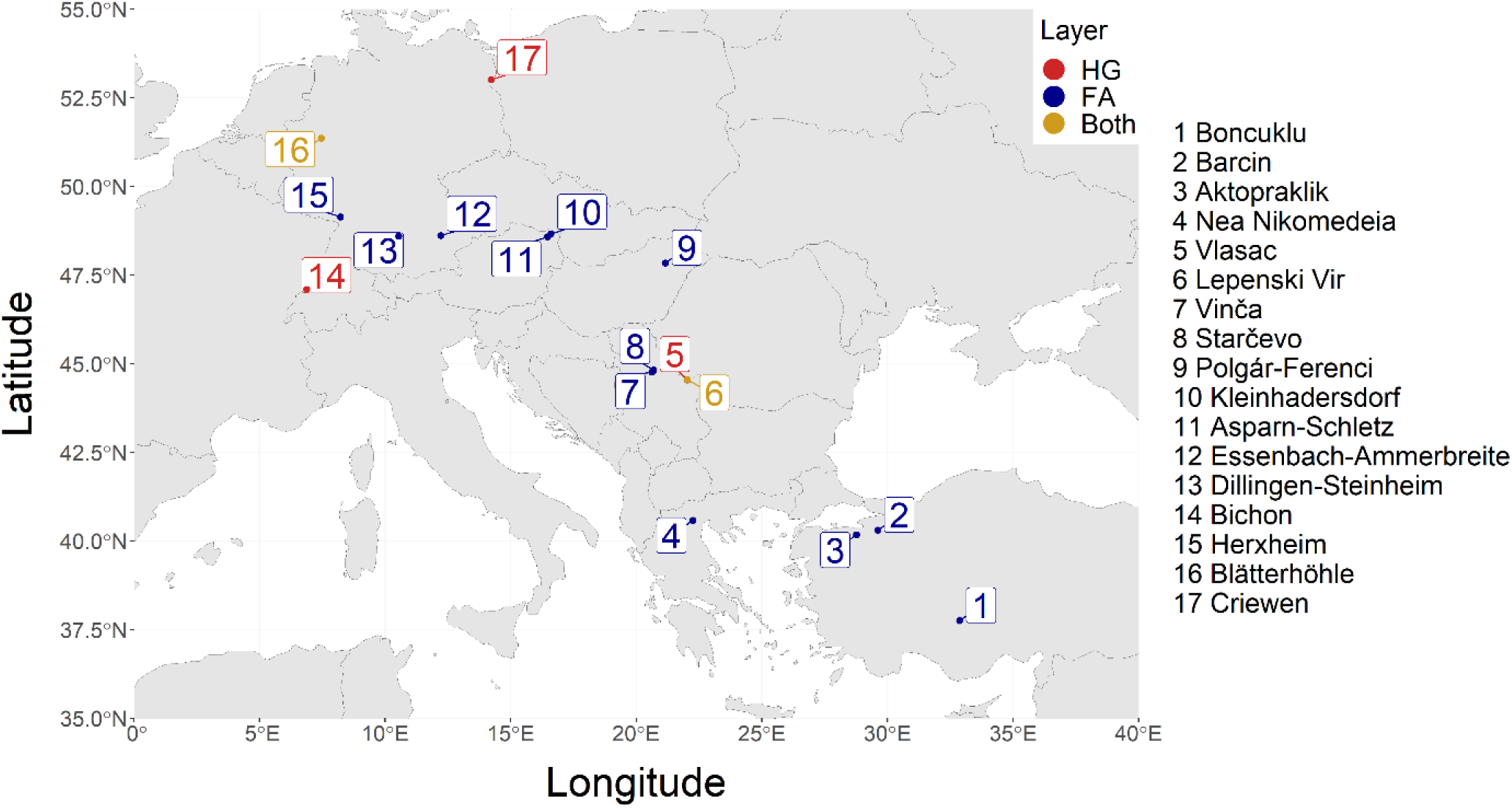
Geographical distribution of the 67 palaeogenomes used in the present study. Locations of hunter-gatherer samples are designated with “HG”, locations of farmer samples are designated with “FA”, while locations from which both hunter-gatherer and farmer samples were used are designated with “Both”.

We used ATLAS (commit=b745621, *66*) to infer genotype likelihoods (task=GLF) for all individuals, excluding three bases near the 3’ and 5’-ends of reads. Subsequently, we computed major and minor alleles (task=majorMinor) from these genotype likelihoods, creating a file that delineates variant call sites and their known alleles. As a measure of the genetic differentiation between the genomes, we used the pairwise pseudo-haploid nucleotide diversity. Thus, to get the pseudo-haploid calls, we used the previously generated file with known alleles and ATLAS (task=call method=majorityBase) to select at each site the base with the most occurrences, filtering out sites with coverage <2x. Then, for each possible pair of genomes, the pairwise nucleotide diversity was estimated with ATLAS (task=estimateF2) as the number of variable sites divided by the number of total sites compared between the two genomes, resulting in 2,211 pairwise values, provided in Supplementary Data S2.

We grouped together genomes that were found in the same geographic location, belonging to the same population group (HG or FA) and with ages differing by less than 350 years, for a total of 23 “population samples”, for the purpose of creating samples of larger size and reducing the total number of pairwise statistics. We estimated 267 mean values of intra-sample and inter-sample pseudo-haploid nucleotide diversity using the estimated 2,211 pairwise values. The mean pseudo-haploid nucleotide diversity within and between samples were used as summary statistics in the next steps of the analysis and are provided in Supplementary Data S3.

### Spatially explicit simulations

To simulate demographic parameters of the Neolithic transition, we used a modified version of the spatially explicit simulator SPLATCHE3 (*67*). SPLATCHE3 uses a map of the studied area in ASCII format, divided into cells, where two overlapping layers of simulated populations, each with its own demographic characteristics. Interactions between populations of each layer may occur according to predefined parameters. Each cell thus contains two demes, one per population, HG or FA. Time is counted in generations of 25 years in this study. The simulations consist of two parts, a forward in time demographic simulation and a backward coalescent genetic simulation, which produces DNA sequences for each genome with user-defined times and location, grouped by samples as in the real dataset. We simulated 1,600 generations with a starting time of the simulations at ∼40,000 YBP, corresponding approximately to the onset of modern human migration into Europe (*68*). The mean estimated ages of the observed genomes (Supplementary Data S1) were transformed into the number of generations since the start of the simulation.

The two simulated layers of demes correspond to HGs and FAs, following previous studies (*29*, *30*) and the map used in the simulations is divided into 75×47 cells of 100×100 *km* and resembles western Eurasia, including the Near East and North Africa (Fig. 1A). Although we focus our analysis on samples distributed along the Continental route, we considered a larger area in the simulations to avoid an arbitrary decision on delimiting this route. Coordinates of the observed dataset were transformed into the WGS84 Plate Carree format with the proj4 R package (*69*). The simulations used two zones of HG and FA competition. The area above latitude 43.2° (Fig. 1B) is termed “Northwest Continental route” hereafter while the area below that latitude is termed “Southeast Continental route”. In each cell, admixture occurs unidirectionally at a rate γ from HGs to FAs using the SPLATCHE3 assimilation model (*67*). This means that at each generation, when HGs and FAs coexist in a cell, a fraction γ of the contact between them results in admixture. This leads to the passage of HG genes into the FA population, and thus to gene flow from HG to FA, following previous simulation studies (*29*, *30*). Note that the model does not distinguish whether this gene flow is due to the adoption of agriculture by HGs, the assimilation of HGs into the agricultural population, or mixed encounters between individuals from the two populations resulting in the birth of a child in the FA population.

In the modified version of SPLATCHE3 used in this study (available at Zenodo.org), some specific features were added, to allow spatiotemporal variation in interactions between the two simulated populations:

a. The admixture rate (*γ*) is considered to vary both spatially and temporally. Each deme is endowed with a value of γ, which is subject to modification at times specified by the user.
b. Temporally increasing admixture rate. *γ* can either be the set value for the whole duration of the simulation or start from 0 and increase each generation by a fraction of the set value, until it reaches that value. The fraction of *γ* by which it increases in each generation is referred to as *γ_inc_*.
c. Two competition zones. The competitive interactions between the two simulated layers are modeled using a Lotka-Volterra model of competition (*70*, *71*). Notably, each pair/group of demes within these zones is assigned a unique competition coefficient (*α*). The utilization of different α for each area allows us to delineate zones where the two simulated populations coexist for varying durations (inversely proportional to the value of α). In all cases, α is considered symmetrical between HGs and FAs in a given zone.
d. Sequencing error. A sequencing error probability (ε) defined by the user has been added to SPLATCHE3, which can now introduce errors to the generated DNA sequences (Supplementary Text 5), to better reproduce observed data.

Parameter values for each simulation were sampled from uniform distributions described in Table 2. The carrying capacity of each deme (parameter *K*) is given as the effective haploid population size. For HGs, the range of possible *K_HG_* investigated was 270-450. The range (density between 0.04 and 0.0675 individuals/*km*^2^) was based on the 95% confidence interval of the estimated final Palaeolithic population in Europe at 13,000 YBP (*72*), multiplying the density by the deme size to get the census population size, dividing it by 3 to get the effective population size of diploid individuals (*53*) and, finally, multiplying it by two to get the effective population size in number of gene copies. The range of *K* for FAs (*K_FA_*) was 1,200-2,400 (0.18 - 0.36 *km*^2^), starting from the range estimated by Silva *et al.* (*30*) and adjusting it according to preliminary simulations. The population density (*N_t_*) in each deme, in each generation *t* is logistically calculated and regulated by *K* and the growth rate parameter (*r*). The chance of an individual in a deme to migrate at each generation is given as the migration rate (*m*). For HGs the values of *m_HG_* and *r_HG_* are 0.15 and 0.2 respectively, following Currat & Excoffier (*29*). For FAs, *r_FA_* was set to 0.55, while *m_FA_* was variable, ranging between 0.1-0.5, which was chosen based on preliminary simulations. The migration can either happen in a stepping-stone manner (*73*), or through Long Distance Dispersal (LDD), with the fraction of migration events that are LDD being an additional parameter (*P_LDD_*). The prior range of *P_LDD_* was set to 0-0.025, based on Rio *et al.* (*32*). The distance the migrants travel is drawn from a gamma distribution, defined by a shape (*β*) and rate (*λ*) parameter, which are equal to 1.209 and 0.15046 respectively (*32*).

**Table 2.** Prior ranges of parameters that were varying between simulations and values of the constant parameters.

| Parameter | Prior distribution |
| --- | --- |
| Admixture rate ($\gamma$) * | 0-0.1 |
| Admixture rate in the Southeast Continental route ($\gamma_S$) †‡ | 0-0.1 |
| Admixture rate in the Northwest Continental route ($\gamma_N$) †‡ | 0-0.1 |
| Number of generations of $\gamma$ increase ($t_{inc}$) †‡ | 1-248 |
| Effective population size of HGs ($K_{HG}$) ‡ | 270-450 |
| Effective population size of FAs ($K_{FA}$) ‡ | 1200-2400 |
| Proportion of Long-Distance Dispersals ($P_{LDD}$) ‡ | 0-0.025 |
| LDD shape parameter ($\beta$) | 1.209 |
| LDD scale parameter ($\theta$) | 0.15046 |
| Coefficient of competition in Northwest Continental route ($\alpha_N$) ‡ | 0.15-0.37 |
| Migration rate of HGs ($m_{HG}$) | 0.15 |
| Migration rate of FAs ($m_{FA}$) ‡ | 0.1-0.5 |
| Growth rate of HGs ($r_{HG}$) | 0.2 |
| Growth rate of FAs ($r_{FA}$) | 0.55 |
| Sequencing Error rate ($\epsilon$) ‡ | 0.000045-0.00012 |

\* only used for scenarios AM1-AM5, † only used for scenario SM6, ‡ estimated with SM6

The competition coefficient between HGs and FAs (*α*) was initially allowed to take on values in the entire possible range (0.0-1.0), then reduced to 0.15-0.37 after a set of exploratory simulations (see Supplementary Text 4) to maximize the number of compatible simulations. Since there were no HGs from the Southern Balkans and western Anatolia included in the dataset, we decided to focus the exploration of the competition coefficient in the “Northwest Continental route”, as defined above, including regions where the HGs cohabitated with the FAs for an extended period of time (*14*). For that purpose, we used the new feature of SPLATCHE3 that allows for two zones of competition (Fig. 1B), and we set the competition *α_N_* in the Northwest Continental route (defined by latitude larger than 43.2°) to be a variable, while in the rest of the map *α_S_* was set to 1, similarly to previous studies (*30*). In addition, we allowed for different admixture rates in each of those two zones to make this parameter variable in space for scenario SM6. HG individuals can move from the HG layer to the FA layer at a rate *γ*, which represents the probability that a contact between HGs and FAs will result in gene flow from HGs to FAs, ranging from 0 to 0.1, based on the estimation by Silva *et al.* (*30*) and adjusted according to preliminary simulations. The sequencing error rate (ε) was set between 0.000045 and 0.00012, with a range chosen from preliminary simulations.

For each genome we simulated 50 loci of 1,000bp each, using a mutation rate of 2.15×10^-8^ mutations per nucleotide site per generation, based on the estimations of Nachman & Crowell (*74*). In the modified version of SPLATCHE3 used, the pseudo-haploid nucleotide diversity within and between samples is calculated directly by the program and written in a separate output file with the extension “.*PseudoHaploDiv*”.

### Five investigated scenarios, with different admixture modes

To investigate whether the rate of admixture between HGs and FAs was uniform along the Continental route and whether it became more frequent the longer their cohabitation went on, we started by elaborating five scenarios, which differed in the spatial and/or temporal mode of admixture (Fig. 1D). These five scenarios (named AM1-AM5) were inspired from those described in a previous study from our group (*30*):

AM1. Constant admixture rate in time and space: the parameter *γ* remains constant across the entire geographical map, maintaining its values from the initial interaction between HGs and FAs to the conclusion of the simulations. This scenario mirrors the demic diffusion model, where a limited number of HGs integrate into the farming community in each deme at a uniform rate over space and time (*29*).

AM2. Spatially increasing admixture rate: in this scenario, the parameter *γ* remains constant over time but exhibits a gradual increase as one moves from Anatolia towards Central Europe. This scenario reflects a variant of the demic diffusion model, including a gradient of admixture between HGs and FAs with higher rates of admixture towards the northwest regions of the Continental route.

AM3. Temporally increasing admixture rate: in this scenario, the parameter *γ* is initially equal to 0 across all demes. However, upon colonization by FAs, *γ* within a deme progressively increases with each succeeding generation at a rate denoted as *γ_inc_*, until it reaches its maximum value equal to *γ* at the end of the cohabitation time with HG. *γ* is constant across all demes. This scenario represents a variant of the demic diffusion model incorporating an escalating admixture rate over time following the introduction of farming into a particular cell (region).

AM4. Spatially and temporally increasing admixture rate: in this combined scenario, the parameter *γ* exhibits a dual pattern of increase: it escalates with distance from Anatolia across geographical space, and it also increases with each successive generation as HGs and FAs coexist within demes. This scenario integrates features from both AM2 (spatial gradient) and AM3 (temporal escalation) reflecting a nuanced demic diffusion model.

AM5. Spatially decreasing admixture rate: in this scenario, the parameter *γ* remains constant over time, but decreases as one moves from Anatolia to Central Europe. Unlike AM2, where there is a spatial gradient of increasing admixture, this scenario represents the opposite trend, with admixture rates decreasing towards the northern regions along the route from Antolia to Central Europe It is important to note that a scenario involving temporally decreasing admixture rates was not explored, as it has been previously dismissed by Lipson *et al*. (*18*).

For scenarios with spatially varying admixture rates (AM2, AM4, AM5), the region between South-West Asia and Central Europe was subdivided into ten zones. Each of these zones was assigned a value for the parameter *γ* that differs (positively or negatively) from the rate in the previous zone by 10% of the maximum *γ*.

To determine the speed *γ_inc_* at which *γ* needs to increase each generation to reach its maximum value by the end of the cohabitation period for scenarios AM3 and AM4, we conducted 200,000 preliminary simulations. These simulations were based on the parameter priors outlined in Table 2 and utilized scenario AM1 as the starting point. From those preliminary simulations, we retrieved the mean cohabitation time between HGs and FAs and used the mgcv R package (*75*) to fit a generalized additive model (GAM), where the mean cohabitation time was regressed against *K_HG_*, *K_FA_* and *α_N_*. The value of *γ_inc_* was then set to the inverse of the mean cohabitation time predicted with the three aforementioned parameters as input, for each simulation of scenarios AM3 and AM4.

We performed 800,000 simulations per scenario (AM1-AM5), of which 33,500 simulations per scenario were retained for further analysis, for a total of 167,500 simulations. This subset refers to simulations that were able to reproduce the observed dataset, meaning those that produced all the observed samples at the appropriate generation, location and population layer (HG or FA), and for which the HGs went extinct before the end of the simulation. Simulations where not all genomes could be sampled (usually because HGs went extinct too early or where HGs survived later than what is assumed), were filtered out before further analysis. We refer to this rejection of simulations as a “demographic filter”. The number of simulations that passed the demographic filter for the various scenarios ranged between 33,500 and 35,000 depending on the scenario. Therefore, if more than 33,500 simulations successfully passed the “demographic filter” for any given scenario, then a subset of 33,500 simulations was drawn randomly to get an equal number of simulations for the model choice procedure. To evaluate the robustness of the scenario comparison, we built a confusion matrix by using ABC random forest, using all retained simulations for each scenario. The five scenarios were first tested by using a simulated dataset with homogeneous genome distribution in space and time, to verify their ability to produce different/meaningful results (see Supplementary Text 1).

### Scenario of the Continental Neolithic expansion for parameter estimation

Additionally, we introduced a sixth scenario (referred to as SM6) that incorporates the characteristics of all previous scenarios. In SM6, the spatial aspect of admixture was integrated by exploiting the same concept of defining two zones as with differing competition, similar to the “Southeast Continental route” and “Northwest Continental route” described earlier. This setup allowed us to test whether one zone exhibited higher values of γ compared to the other (indicating spatially increasing or decreasing γ) or if both zones maintained the same γ values (indicating γ could be considered constant in space).

For the temporal heterogeneity of γ, we randomly chose for each simulation a number of generations, ranging between 1 and the mean cohabitation time of HGs and FAs, and we set the value of the *γ_inc_* to the inverse of the chosen number of generations. That way γ could be constant in time (if the chosen number of generations was 1) or increase for a flexible time period, up to the maximum mean number of generations of cohabitation. The number of generations during which the γ increased ranged from 1 to 248. However, it did not follow a uniform distribution, as its values depended on the estimated mean cohabitation time for each simulation. It is worth noting the following: while γ was increasing steadily with each passing generation until reaching its maximum value at the end of the cohabitation period in scenarios AM1-AM5, in scenario SM6 it was set to increase for a specific number of generations and then retain its maximum value until the end of the cohabitation period. For SM6, we performed 2,400,000 simulations, 100,000 of which were retained after applying the demographic filter.

The average distance of the LDD events in our simulations was 800 km. To test if shorter LDD events affect the resulting levels of pseudo-haploid nucleotide diversity, we performed a set of simulations using scenario SM6 and an average LDD distance of 300 km. Additionally, we performed a set of simulations where all FA migrations were short scale (*P_LDD_* = 0), to investigate the importance of LDDs in replicating the observed dataset.

### Statistical Analysis: ABC Model choice and parameter estimation

The choice of the most likely scenario was performed with the abcrf R package (*76*), which uses a random forest approach to perform the ABC selection. The choice of abcrf was motivated by its superior discriminative power even with a lower number of simulations compared to other ABC methods. Furthermore, abcrf is capable of handling a large number of summary statistics effectively. In our study, we utilized 267 summary statistics of mean pseudo-haploid nucleotide diversity within and between samples, leveraging the robust capabilities of abcrf for our analyses (see Supplementary Text 1). The random forest ABC analysis outputs the number of trees that selected each model and the posterior probability of the best model. The forest consisted of 2,000 trees after checking that a larger number of trees does not significantly decrease the scenario misclassification rate.

Given that abcrf does not formally evaluate the fit of the simulated data to the observed data, we assessed the probability of each of the five scenarios reproducing the observed data using ABCtoolbox2 (*77*). This assessment involved calculating the marginal density p-value, where the null hypothesis posits that the simulated data fit the observed values adequately. Due to the high number of summary statistics, we used the 10 first components that best explain the variability of the data for estimating the marginal density p-value. The components resulted from the computation of linear combinations of the 267 values of intra- and inter sample pseudo-haploid nucleotide diversity using Partial Least Squares (PLS, *78*). This transformation was done with the find “pls.R” R script and the “transformer” software (both included in ABCtoolbox2), to reduce the dimensionality of the dataset. The marginal density is estimated on retained simulations based on a tolerance level (δ) and the Euclidean distance between each simulated and the observed summary statistics. For scenarios AM1-AM5, the parameter δ was fixed at 0.03, and we retained 1,000 simulations per scenario. Two additional values of δ were tested for scenarios AM1-AM5, and the results are presented in Table S1. For scenario SM6, three values of the parameter δ were used (0.005, 0.01, and 0.05), with 500, 1,000, and 5,000 simulations retained respectively.

We made demographic inferences by estimating the most probable values of the parameters used in the simulations with scenario SM6 (‡ in Table 2). The parameter estimation was performed with ABCtoolbox2, which calculates the minimized Euclidean distance between simulated and observed statistics. Based on these distances, ABCtoolbox2 retains a fraction of the simulations that are closer to the observed data and computes the posterior distribution of the parameter values (*77*). This is done using an ABC-GLM regression adjustment (*79*). We performed the parameter estimation using three values of δ, 0.005, 0.01 and 0.05, retaining 500, 1000 and 5000 simulations respectively. The results for δ=0.01 are presented in the main text, while the results of the other values of δ are presented in Table S3. For the two γ variables (γ*_S_* and γ*_N_*) and the two K variables (K*_HG_* and K*_FA_*), we estimated two-dimensional posteriors and used a MCMC to sample from these posteriors, in order to investigate if there are combinations of these variables that are more probable than others.

The summary statistics used for the parameter estimation were the 10 first components resulting from the PLS transformation of the pseudo-haploid nucleotide diversity. For each parameter we estimated the posterior range, as the 95 High Density Interval (HDI) of the distribution, and we considered its mode and mean as point estimates. We evaluated the precision of our point estimates by calculating their 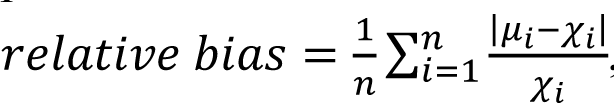, where *n* is the number of pseudo-observations, *χ_i_* is the _𝑛_ 𝑖=1 𝜒_𝑖_ pseudo-observed parameter value, and *μ_i_* is the mean or mode of the posterior parameter distribution of the replicate. To do this, we considered the summary statistics of each retained simulation, in turn, as pseudo-observed statistics with known parameter values and we performed the parameter estimation procedure with all other simulations.

## Author contributions

MC conceived and coordinated the study. AT, MC, JBu, CSQ and DW designed the study. JBl contributed to the dataset. CSRB processed the dataset. AT did all the analyses and simulations with the help of MC, CSQ and DW. AT, MC, MB, JBu and CSQ interpreted the results. AT wrote the first draft. All authors contributed to the writing of the manuscript and agreed on the final version.

## Data and materials availability

All data are available in the main text or the supplementary materials. All genomes analyzed in this study were already published and references are provided in Supplementary Data S1. The modified version of SPLATCHE3 is provided in the online free repository Zenodo.org (link: https://zenodo.org/records/10495323).

