## Supplementary Materials for "Local increases in admixture with hunter-gatherers followed the initial expansion of Neolithic farmers across continental Europe"

**This PDF file includes:**

Supplementary Text 1-5

Figs. S1 to S7

Tables S1 to S8

**Table S1. Marginal posterior density p-values of the five investigated scenarios.** Calculation performed with ABCtoolbox2 (*77*), using 33,500 simulations per scenario and three separate tolerance levels, retaining 500, 1000 and 1,675 simulations (tolerance of 0.015, 0.03 and 0.05 respectively). For this analysis, the observed values of pseudo-haploid nucleotide diversity were transformed into PLS components.

| | Marginal posterior density p-value with tolerance 0.015 | Marginal posterior density p-value with tolerance 0.03 | Marginal posterior density p-value with tolerance 0.05 |
|---|---|---|---|
| AM1 - Constant assimilation rate in space and time | 0.45 | 0.31 | 0.91 |
| AM2 - Spatially increasing assimilation rate | 0.12 | 0.65 | 1.00 |
| AM3 - Temporally increasing assimilation rate | 0.46 | 0.89 | 0.28 |
| AM4 - Spatially and temporally increasing assimilation rate | 0.39 | 0.67 | 0.46 |
| AM5 - Spatially decreasing assimilation rate | 1.00 | 0.24 | 0.82 |

**Table S2. Model choice between the five investigated scenarios.** Performed with abcrf R package (random forest approach, *76*), using 2,000 trees and 33,500 simulations per scenario. The votes refer to the number of trees that selected each scenario as the most probable one. For the analysis, the untransformed summary statistics were used.

| AM1 Constant assimilation rate in space and time votes | AM2 Spatially increasing assimilation rate votes | AM3 Temporally increasing assimilation rate votes | AM4 Spatially and temporally increasing assimilation rate votes | AM5 Spatially decreasing assimilation rate | Chosen scenario | Posterior probability of chosen scenario |
|---|---|---|---|---|---|---|
| 363 | 403 | 446 | 430 | 358 | AM3 | 0.43 |

**Table S3. Characteristics of the estimated parameters, using the simulations of scenario SM6.** The tolerance levels of 0.005, 0.01 and 0.05 correspond to 500, 1,000 and 5,000 retained simulations out of the 100,000 simulations of the scenario. The estimation was performed with ABCtoolbox2 (*77*).

| Parameter | Tolerance level | Prior minimum | Prior maximum | Posterior Mode | Posterior Mode bias | Posterior Mean | Posterior Mean bias | Posterior Lower HDI95 | Posterior Upper HDI95 |
|---|---|---|---|---|---|---|---|---|---|
| Assimilation rate in Southern Continental route ($\gamma_S$) | 0.005 | 0 | 0.1 | 0.029 | 4.64 | 0.047 | 4.13 | 0.002 | 0.092 |
| | 0.01 | 0 | 0.1 | 0.024 | 3.96 | 0.049 | 3.67 | 0.003 | 0.096 |
| | 0.05 | 0 | 0.1 | 0.029 | 3.33 | 0.049 | 3.19 | 0.004 | 0.096 |
| Assimilation rate in Northern Continental route ($\gamma_N$) | 0.005 | 0 | 0.1 | 0.035 | 3.25 | 0.049 | 2.73 | 0.004 | 0.096 |
| | 0.01 | 0 | 0.1 | 0.029 | 4.38 | 0.046 | 3.77 | 0.002 | 0.093 |
| | 0.05 | 0 | 0.1 | 0.041 | 2.79 | 0.049 | 3.09 | 0.003 | 0.094 |
| Number of generations of $\gamma$ increase ($t_{inc}$) | 0.005 | 1 | 248 | 60 | 1.35 | 72 | 2.18 | 3 | 137 |
| | 0.01 | 1 | 248 | 70 | 1.49 | 72 | 2.29 | 2 | 136 |
| | 0.05 | 1 | 248 | 55 | 1.4 | 65 | 2.2 | 1 | 129 |
| Effective population size of HGs ($K_{HG}$) | 0.005 | 270 | 450 | 335 | 0.13 | 355 | 0.11 | 275 | 433 |
| | 0.01 | 270 | 450 | 343 | 0.14 | 357 | 0.12 | 273 | 435 |
| | 0.05 | 270 | 450 | 337 | 0.14 | 354 | 0.12 | 272 | 435 |
| Effective population size of FAs ($K_{FA}$) | 0.005 | 1200 | 2400 | 1430 | 0.19 | 1763 | 0.17 | 1230 | 2332 |
| | 0.01 | 1200 | 2400 | 1479 | 0.18 | 1769 | 0.17 | 1230 | 2322 |
| | 0.05 | 1200 | 2400 | 1467 | 0.18 | 1736 | 0.17 | 1200 | 2269 |
| Proportion of Long-Distance Dispersals ($P_{LDD}$) | 0.005 | 0 | 0.025 | 0.021 | 2.87 | 0.014 | 4.27 | 0.002 | 0.025 |
| | 0.01 | 0 | 0.025 | 0.019 | 2.26 | 0.013 | 3.50 | 0.002 | 0.024 |
| | 0.05 | 0 | 0.025 | 0.016 | 1.81 | 0.013 | 2.63 | 0.002 | 0.024 |
| Coefficient of competition in Northern Continental route ($\alpha_N$) | 0.005 | 0.15 | 0.35 | 0.214 | 0.17 | 0.24 | 0.19 | 0.15 | 0.336 |
| | 0.01 | 0.15 | 0.35 | 0.203 | 0.17 | 0.231 | 0.19 | 0.15 | 0.327 |
| | 0.05 | 0.15 | 0.35 | 0.199 | 0.18 | 0.227 | 0.19 | 0.15 | 0.321 |
| Migration rate of FAs ($m_{FA}$) | 0.005 | 0.1 | 0.5 | 0.411 | 0.38 | 0.340 | 0.36 | 0.163 | 0.5 |
| | 0.01 | 0.1 | 0.5 | 0.423 | 0.40 | 0.344 | 0.37 | 0.162 | 0.5 |
| | 0.05 | 0.1 | 0.5 | 0.427 | 0.41 | 0.339 | 0.4 | 0.153 | 0.5 |
| Sequencing Error rate ($\varepsilon$) | 0.005 | 4.50E-05 | 0.00012 | 0.000059 | 0.216 | 0.000074 | 0.222 | 0.000045 | 0.000107 |
| | 0.01 | 4.50E-05 | 0.00012 | 0.000057 | 0.217 | 0.000071 | 0.224 | 0.000045 | 0.000103 |
| | 0.05 | 4.50E-05 | 0.00012 | 0.000056 | 0.207 | 0.000071 | 0.216 | 0.000045 | 0.000102 |

**Supplementary Text 1. Theoretical investigation of the five scenarios**

Objective: In order to check whether the five demographic scenarios we have studied in the main text are capable of producing identifiable results and whether the random forest ABC (*76*) has enough discriminatory power, even at low numbers of simulations, we performed a set of exploratory simulations with SPLATCHE3.

The five investigated scenarios (AM1 to AM5) are the same as in the main text, see Material and Methods for details about the parameters. The only difference is the molecular dataset we used. Here we used an “ideal” dataset homogeneous in space and time. The sampled dataset consisted of both HGs and FAs in five locations and at five different times, for a total of 50 samples. The sampling locations were in equidistant demes (Fig. S1). The first sampling date was shortly after the colonization of the entire map by FAs (generation 1,250), and we sampled in increments of 30 generations (~750 years). The parameter priors were the same as in Table 2 of the main text.

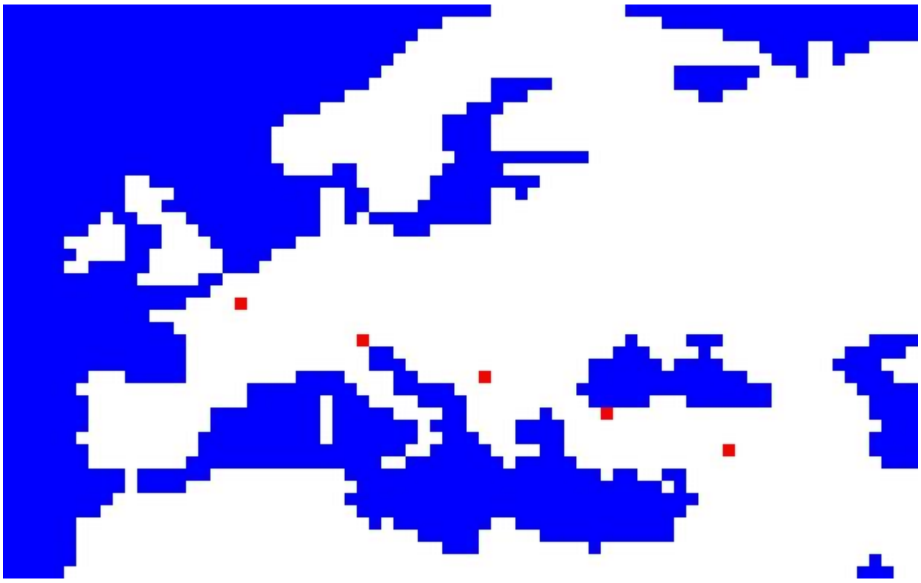

**Fig. S1. Sampling locations for the simulations investigating the differences between scenarios.**

We performed 25,000 simulations per scenario. 13,000 simulations were retained from each scenario, after filtering out the ones missing samples (demographic filter). Contrary to the main text, we retained the ones where the HGs existed until the end of the simulation, since our goal was a theoretical investigation of the scenarios and not to replicate the observed demographic history of the Neolithic.

We constructed a confusion matrix with abcrf, using as summary statistics all the pairwise values of inter- and intra-sample pseudo-haploid nucleotide diversity (Fig. S2), as in the main text analysis.

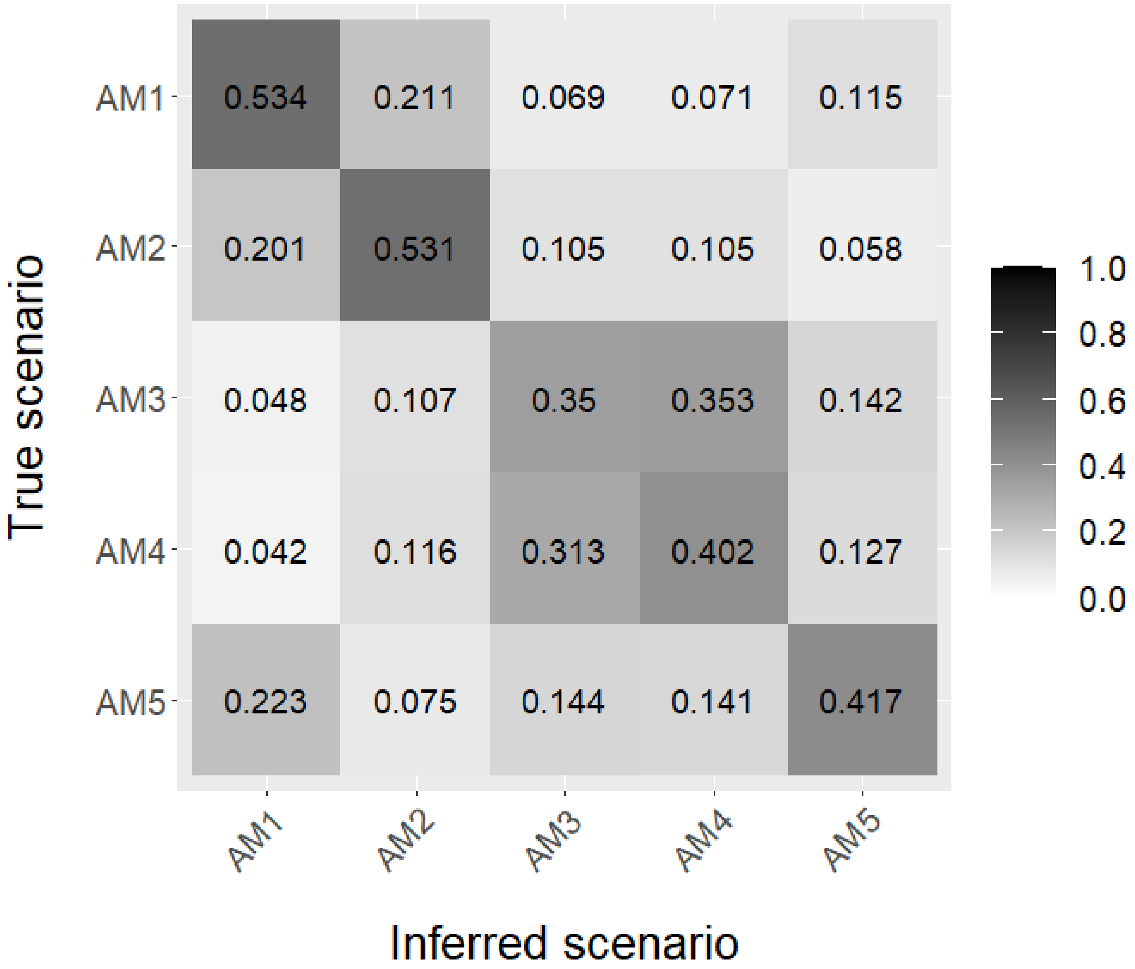


**Fig. S2. Confusion matrix of the five investigated scenarios using pseudo-haploid nucleotide diversity.** Calculation performed with abcrf R package (random forest approach, *76*), using 13,000 simulations per scenario and 2,000 trees. On the first column is given the actual scenario and on the first row the predicted scenario, while the values correspond to the relative frequency with which a scenario was identified as each of the predicted scenarios.

The model choice validation with abcrf using an artificial dataset homogeneous in space and time shows variation in the accuracy of scenario identification when using all the pairwise values of pseudo-haploid nucleotide diversity as summary statistics. The misclassification rate is 0.55, with scenarios AM1 and AM2 having a probability of ~0.55 to be correctly identified, which means the constant assimilation rate and the spatially increasing one create an identifiable pattern of nucleotide diversity. The two scenarios with temporally increasing assimilation rate (AM3 and AM4) were hardly distinguishable between them, but together were clearly differentiated from the other three scenarios of constant assimilation rate (AM1, AM2 and AM5), which means that the temporal increase of the assimilation rate creates a distinct pattern of nucleotide diversity, but also erases the spatial pattern. Despite the relatively high misclassification rate, there exists a detectable signal that differentiates the simulated scenarios with the information lost resulting from the pseudo-haploidization of the genomic data.

Additionally, we tested if using a PLS transformation on the data (see main text), improves the ability of abcrf to correctly identify the scenarios. We transformed the pairwise values of inter- and intra-sample pseudo-haploid nucleotide diversity into PLS components for all five scenarios and we used the first 10 components and abcrfr to construct a confusion matrix (Fig. S3). It shows that a PLS transformation does not improve the ability of ABC to identify the various scenarios. For this reason, we chose to use the original pairwise values of inter- and intra-sample pseudo-haploid nucleotide diversity as summary statistics for the scenario choice on the observed data.

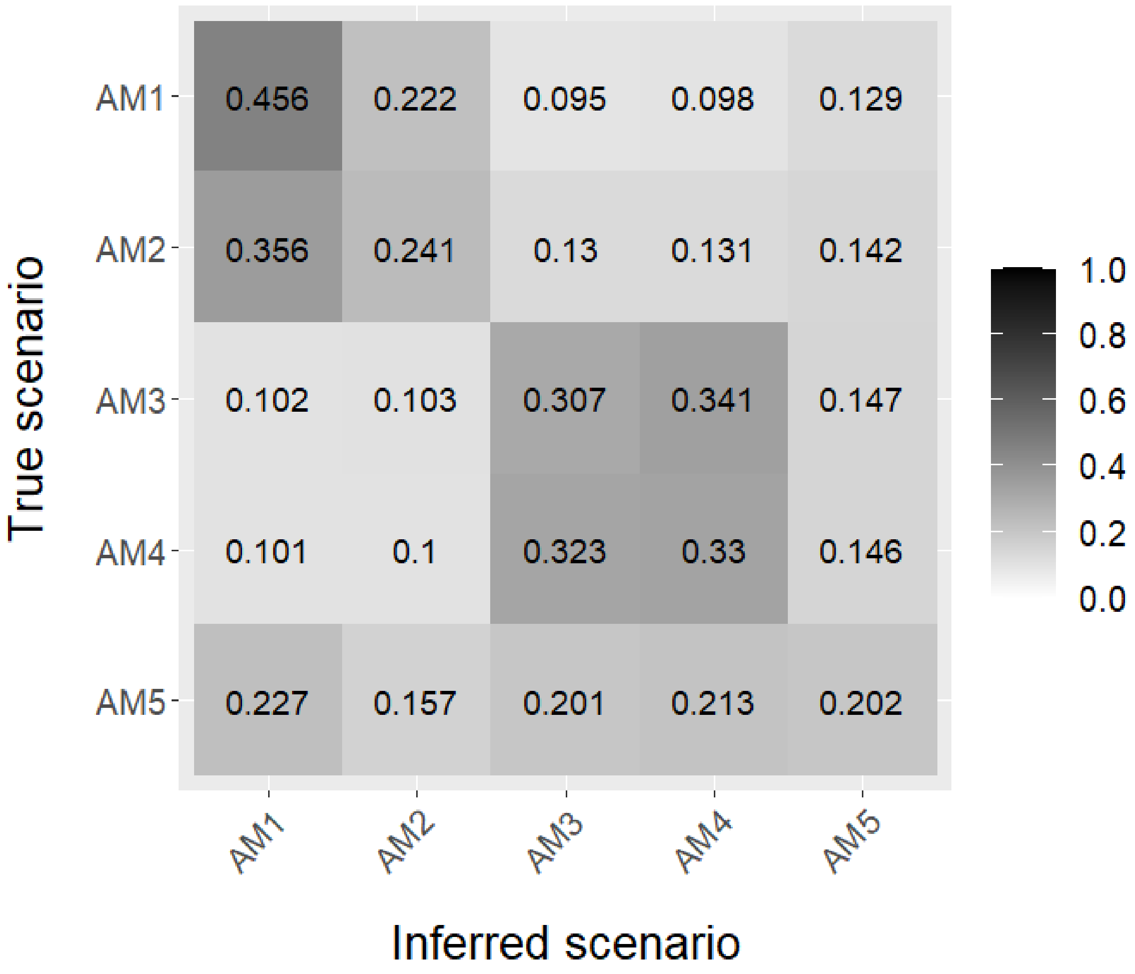


**Fig. S3. Confusion matrix of the five investigated scenarios using PLS transformed pairwise values of inter- and intra-sample pseudo-haploid nucleotide diversity.** Calculation performed with abcrf R package (random forest approach, *76*), using 13,000 simulations per scenario and 2,000 trees. On the first column is given the actual scenario and on the first row the predicted scenario, while the values correspond to the relative frequency with which a scenario was identified as each of the predicted scenarios.

**Supplementary Text 2. Model choice and parameter estimation omitting genomes from Lepenski Vir**

Objective: Lepenski Vir is a special location in the history of the European Neolithic, and a categorization of genomes from this location as hunter-gatherers (HG) or farmers (FA) is challenging (*19*). To remove the uncertainty of the classification from our analysis, we performed the model choice and the parameter estimation after removing the Lepenski Vir samples from both the observed and the simulated summary statistics. Of the original 67 genomes, 59 were retained for this analysis.

*S2.1 Model choice using scenarios AM1-AM5*

All five investigated scenarios were able to reproduce the observed summary statistics, after removing the Lepenski Vir genomes (Table S4).

**Table S4. Marginal posterior density p-values of the five investigated scenarios without genomes from Lepenski Vir.** Calculation performed with ABCtoolbox2 (*77*), using 33,500 simulations per scenario and a tolerance level of 0.03, retaining 1,000 simulations. For this analysis the observed values of pseudo-haploid nucleotide diversity were transformed into PLS components.

| | Marginal posterior density p-value with tolerance 0.03 |
|---|---|
| AM1 - Constant assimilation rate in space and time | 0.90 |
| AM2 - Spatially increasing assimilation rate | 0.99 |
| AM3 - Temporally increasing assimilation rate | 0.86 |
| AM4 - Spatially and temporally increasing assimilation rate | 0.68 |
| AM5 - Spatially decreasing assimilation rate | 0.55 |

The model choice showed that scenario AM3 was still the most probable (Table S5), with its posterior probability (40%) being very close to the one estimated when including the Lepenski Vir genomes (43%).

**Table S5. Model choice between the five investigated scenarios without the genomes from Lepenski Vir.** Performed with abcrf R package (random forest approach, *76*), using 2,000 trees and 33,500 simulations per scenario. The votes refer to the number of trees that selected each scenario as the most probable one. For the analysis, the untransformed pairwise values of inter- and intra-sample pseudo-haploid nucleotide diversity were used.

| AM1 Constant assimilation rate in space and time votes | AM2 Spatially increasing assimilation rate votes | AM3 Temporally increasing assimilation rate votes | AM4 Spatially and temporally increasing assimilation rate votes | AM5 Spatially decreasing assimilation rate | Chosen scenario | Posterior probability of chosen scenario |
|---|---|---|---|---|---|---|
| 340 | 381 | 452 | 426 | 401 | AM3 | 0.40 |

The confusion matrix (Fig. S4) is very similar to the one based on summary statistics that included Lepenski Vir genomes (Fig. 2 of the main text), which shows that the samples from Lepenski Vir did not confound the scenario differentiation.

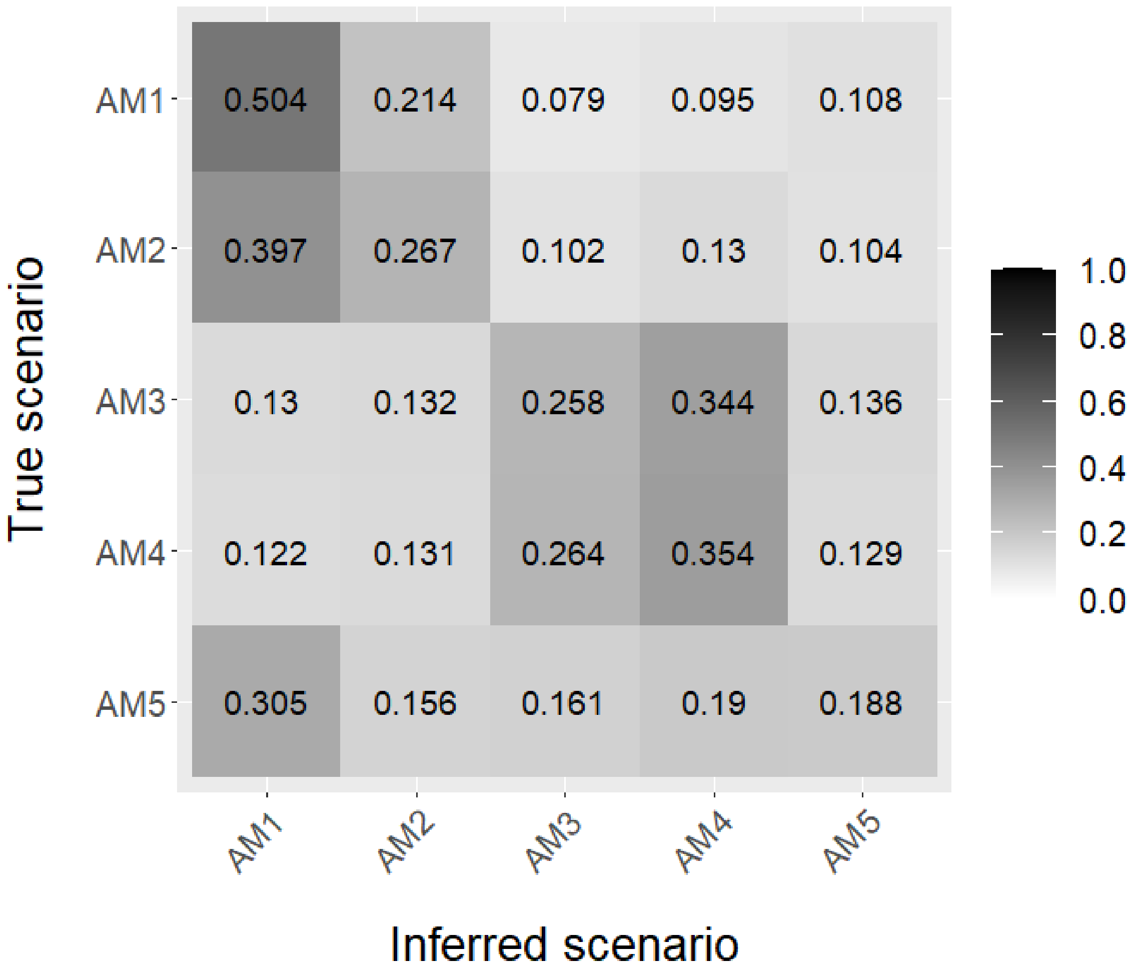


**Fig. S4. Graphical representation of the confusion matrix for three investigated scenarios without the genomes from Lepenski Vir.** Calculation performed with abcrf R package (random forest approach, *76*), using 33,500 simulations per scenario and 2,000 trees.

For the analysis, the untransformed pairwise values of inter- and intra-sample pseudo-haploid nucleotide diversity were used.

*S2.2 Parameter estimation using scenario SM6*

As in the main text, the parameter estimation was performed using scenario SM6. The marginal posterior density p-value of the scenario was 0.48, given a tolerance of 0.01. The characteristics of the posterior distributions of most parameters (except the assimilation rates, TableS6 and Fig. S5) are quite similar to the estimations using the Lepenski Vir genomes. The only important difference is found in the mode of the posterior distribution of the assimilation rates ($\gamma_S$ and $\gamma_N$), as the modes without the Lepenski Vir (7% and 8.3%, respectively) are approximately two times larger than the modes with Lepenski Vir (2.4% and 2.8%). However, both estimations (with and without Lepenski Vir) are characterized by closer means (4.9% and 4.6% *vs* 5.2% and 5.2%), large relative posterior biases and overlapping 95HDI. Therefore we cannot safely conclude that admixture rates estimated with and without the genomes from Lepenski Vir are actually different.

**Table S6. Characteristics of the estimated parameters, using the simulations of scenario SM6 without the genomes from Lepenski Vir.** A tolerance level of 0.01 was used, retaining 1,000 simulations out of the 100,000 simulations of the scenario. The estimation was performed with ABCtoolbox2 (*77*).

| Parameter | Posterior Mode | Posterior Mode bias | Posterior Mean | Posterior Mean bias | Posterior Lower HDI95 | Posterior Upper HDI95 |
|---|---|---|---|---|---|---|
| Assimilation rate in Southern Continental route ($\gamma_S$) | 0.07 | 3.78 | 0.052 | 3.66 | 0.006 | 0.1 |
| Assimilation rate in Northern Continental route ($\gamma_N$) | 0.0838 | 4.46 | 0.056 | 3.04 | 0.01 | 0.1 |
| Number of generations of $\gamma$ increase ($t_{inc}$) | 75 | 1.27 | 75 | 1.85 | 5 | 138 |
| Effective population size of HGs ($K_{HG}$) | 414 | 0.14 | 371 | 0.11 | 290 | 449 |
| Effective population size of FAs ($K_{FA}$) | 1467 | 0.18 | 1710 | 0.17 | 1200 | 2248 |
| Proportion of Long-Distance Dispersals ($P_{LDD}$) | 0.015 | 2.4 | 0.013 | 3.97 | 0.001 | 0.024 |
| Coefficient of competition in Northern Continental route ($\alpha_N$) | 0.212 | 0.18 | 0.238 | 0.19 | 0.15 | 0.3284 |
| Migration rate of FAs ($m_{FA}$) | 0.415 | 0.42 | 0.315 | 0.41 | 0.13 | 0.496 |
| Sequencing Error rate ($\varepsilon$) | 0.00006 | 0.23 | 0.00008 | 0.22 | 0.000055 | 0.00011 |

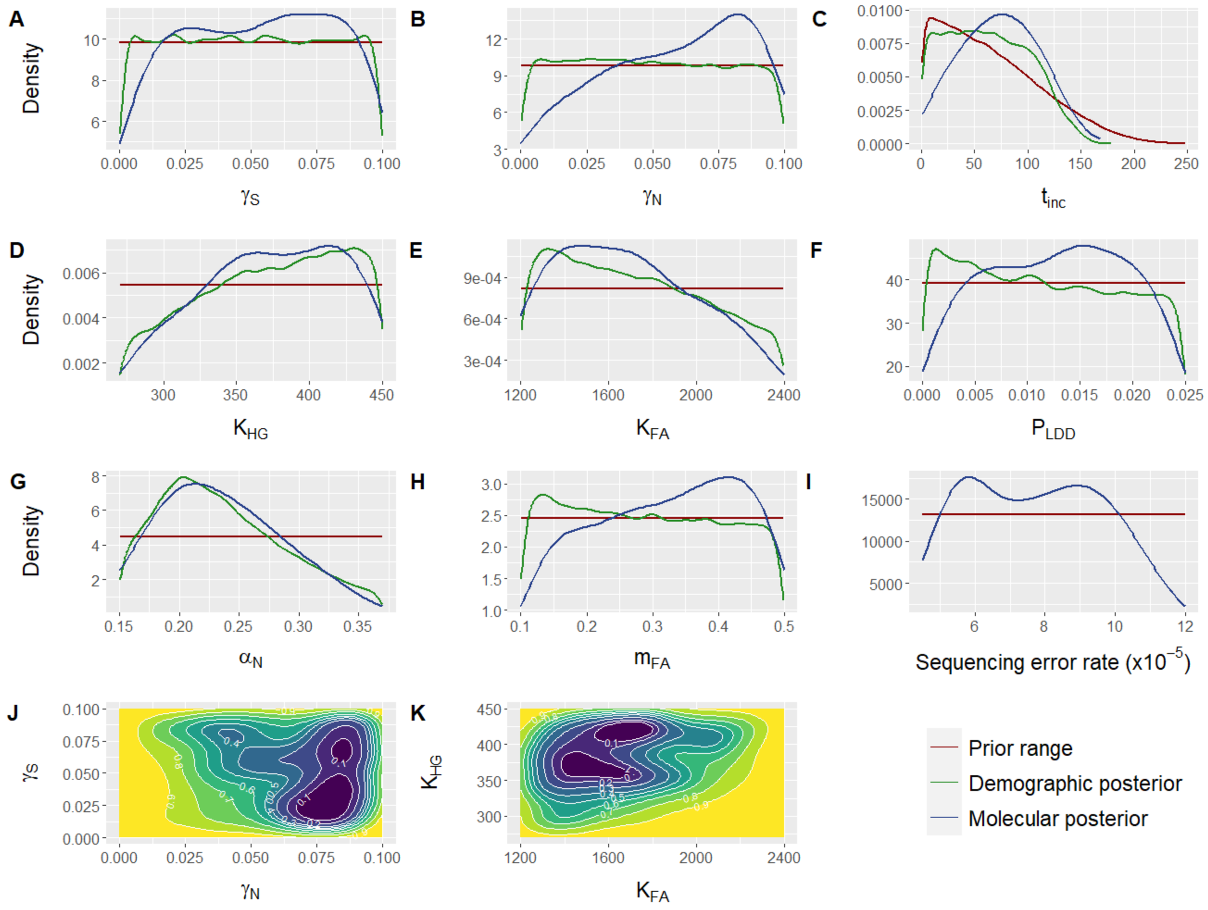


**Fig. S5. Distribution of the estimated parameter values without the genomes from Lepenski Vir.** In panels A-I, the red line corresponds to the prior distribution, the green one to the distribution of parameter values of the simulations that were able to reproduce the observed dataset (demographic posterior), and the blue line corresponds to the posterior resulting from the ABC estimation (molecular posterior). **(A)** Assimilation rate along the Southern Continental route ($\gamma_S$); **(B)** Assimilation rate along the Northern Continental route ($\gamma_N$); **(C)** Number of generations during which the assimilation rate increases ($t_{inc}$); **(D)** Effective population size of HGs ($K_{HG}$); **(E)** Effective population size of FAs ($K_{FA}$); **(F)** Proportion of FA migrations that are Long Distance Dispersals ($P_{LDD}$); **(G)** Competition coefficient along the Northern Continental route ($\alpha_N$); **(H)** Migration rate of FAs ($m_{FA}$); **(I)** Sequencing error rate (ε); **(J)** Two-dimensional posterior distribution of assimilation rate along the Southern Continental route ($\gamma_S$) against the assimilation rate along the Northern Continental route ($\gamma_N$); **(K)** Two-dimensional posterior distribution of effective population size of HGs ($K_{HG}$) against the Effective population size of FAs ($K_{FA}$).

The above analyses show that the uncertainty in group classification of the genomes from Lepenski Vir does not substantially affect the results and the estimations performed in the main text.

**Supplementary Text 3. Model choice using three of the investigated scenarios**

Objective: Since the comparison of the five scenarios showed that both scenarios with temporally increasing assimilation rate (AM3 and AM4) are hardly distinguishable, while the scenario with assimilation decreasing with space (AM5) is almost unidentifiable, we decided to perform again the model choice using only three of the scenarios to avoid noise: AM1 as reference with admixture rate constant in time and space, AM2 for investigating a spatial pattern in the assimilation rate and AM3 for investigating a temporal pattern. Model choice shows again that the most probable scenario is the one with temporally increasing assimilation rate (Table S7).

**Table S7. Model choice performed on three of the investigated scenarios.** Performed with abcrf R package (random forest approach, *76*), using 2,000 trees and 33,500 simulations per scenario. The votes refer to the number of trees that selected each scenario as the most probable one. For the analysis, the untransformed pairwise values of inter- and intra-sample pseudo-haploid nucleotide diversity were used.

| AM1 - Constant assimilation rate in space and time votes | AM2 - Spatially increasing assimilation rate votes | AM3 - Temporally increasing assimilation rate votes | Chosen scenario | Posterior probability of chosen scenario |
|---|---|---|---|---|
| 630 | 660 | 710 | AM3 | 0.52 |

The confusion matrix (Fig. S6) shows that, while assimilation increasing with time produces different results than when it is constant, the spatial heterogeneity of admixture can hardly be distinguished from a scenario of admixture being constant in space.

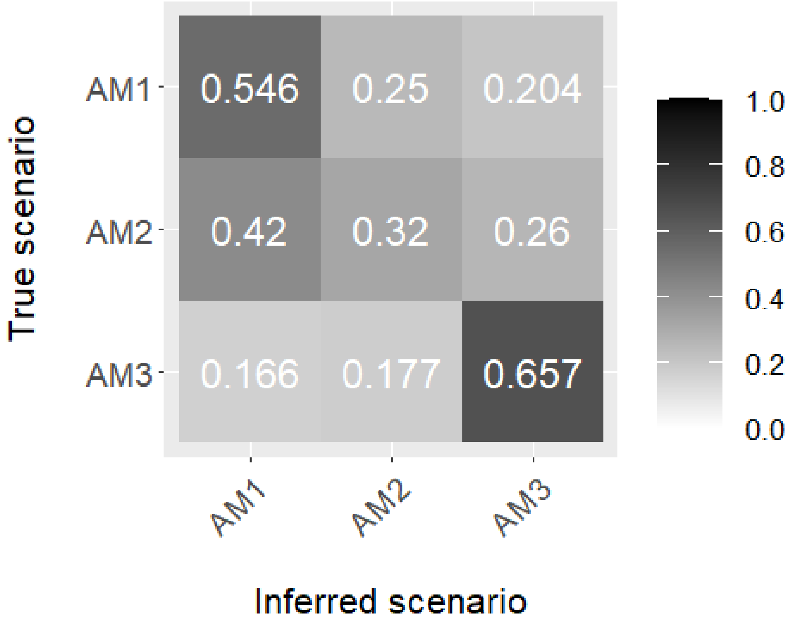


**Fig. S6. Graphical representation of the confusion matrix for three investigated scenarios.** Calculation performed with abcrf R package (random forest approach, *76*), using 33,500 simulations per scenario and 2,000 trees. For the analysis, the untransformed pairwise values of inter- and intra-sample pseudo-haploid nucleotide diversity were used.

**Supplementary Text 4. Exploration of the range of competition coefficient**

Objective: We started by exploring the whole range of possible values of the competition coefficient $\alpha_N$ in the Lotka-Volterra model of competition between HGs and FAs along the Continental route of the Neolithic Transition. The goal was to reduce the prior distribution in order to optimize the computational efficiency.

We performed 25,000 simulations using the modified version of SPLATCHE3 (*67*) described in the main text, allowing for a variable coefficient of competition $\alpha_N$. We followed the same framework as for the simulations of scenario SM6 in the main text. The parameter priors were the same as in Table 2 of the main text, except for $\alpha_N$, for which we explored the whole range of possible values, between 0 and 1. 151 of the simulations (0.6%) passed the demographic filter by successfully reproducing the observed dataset. The posterior range of the competition coefficient is 0.13-0.39, with a 95% High Density Interval (HDI95) of 0.15-0.37. The distribution of values can be seen in Fig. S7.

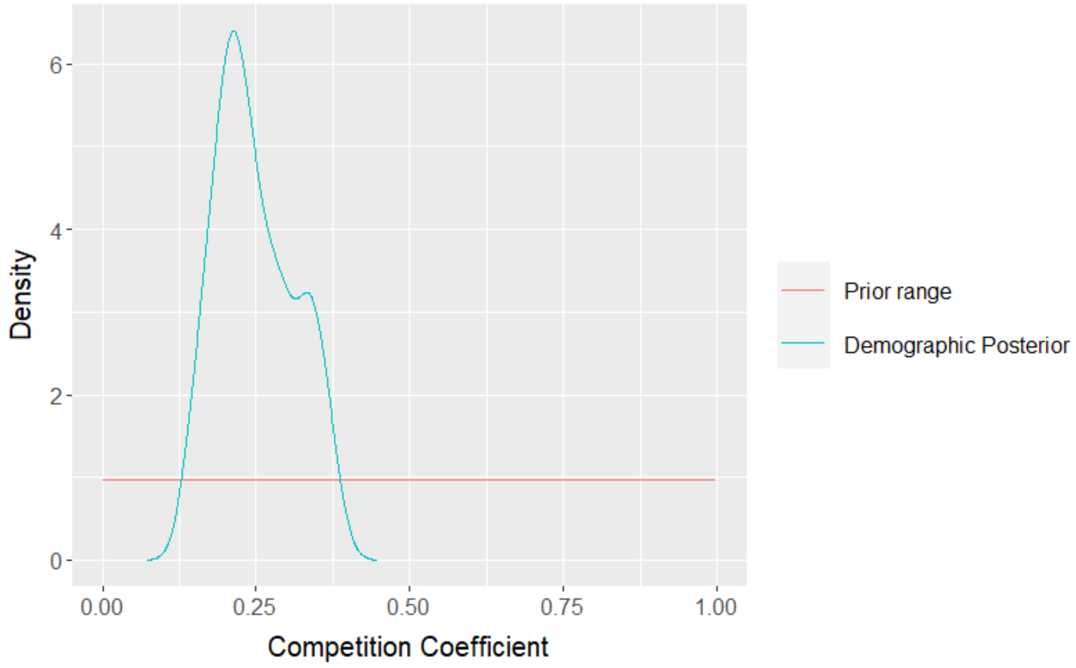


**Fig. S7. Posterior distribution of the values of the competition coefficient.** The red line corresponds to the prior distribution of values used for the simulations, while the light blue corresponds to the distribution of the values of the simulations that were able to reproduce the observed dataset.

Our results show that very low or high values of competition coefficient lead to simulations unable to produce the observed dataset. Therefore, in the following simulations, we reduced the prior range of $\alpha_N$ to 0.15-0.37 to optimize the computational time by avoiding simulating unsuccessful simulations (i.e., not allowing to produce all samples).

**Supplementary Text 5. Implementation of reads sequencing error in SPLATCHE3**

Objective: SPLATCHE3 is not producing sequencing reads but full DNA sequences without errors, so a fixed error rates was implemented to mimic reads errors for pseudo-haploid genomic data as follow:

Table S8 gives the probability to find no difference $P_{nodiff}$ between two genomes (whether intra or inter population samples) at the pseudo-haploid level using the majority allele, given a sequencing error rate ε, which will be set as parameter for SPLATCHE3, and assuming a depth >2.

**Table S8. Equation used to compute the probability $P_{nodiff}$ to find no difference at the pseudo-haploid level between two genomes with known diploid state, given a sequencing error rate ε (using the majority allele and assuming a depth >2).**

| Diploid state | 00 | 01 | 11 |
|---|---|---|---|
| 00 | $P_{nodiff}= \varepsilon^2 + (1-\varepsilon)^2$ | $P_{nodiff}$=1/2 | $P_{nodiff}=2\varepsilon(1-\varepsilon)$ |
| 01 | $P_{nodiff}$=1/2 | $P_{nodiff}$=1/2 | $P_{nodiff}$=1/2 |
| 11 | $P_{nodiff}= 2\varepsilon(1-\varepsilon)$ | $P_{nodiff}$=1/2 | $P_{nodiff}=\varepsilon^2 + (1-\varepsilon)^2$ |

If both genotypes are identical (00x00 or 11x11), then one allele is taken at random for each pseudo haploid genome. The probability that they do not differ is either that none has a sequencing error, with a probability $(1-\varepsilon) * (1-\varepsilon) = (1-\varepsilon)^2$ , or both have a sequencing error with a probability equal to $\varepsilon * \varepsilon = \varepsilon^2$. If only one allele has a sequencing error, then both pseudo-haploid genomes will be identified as different.

If both genotypes are homozygous but for a different allele (00x11 or 11x00), then one allele is taken at random for each pseudo haploid genome. The probability that they do not differ results from one of the two alleles having a sequencing error but not the other, with a probability equal $\varepsilon(1-\varepsilon)$, or the reverse, leading to $2\varepsilon(1-\varepsilon)$. If both alleles have a sequencing error, or both have no sequencing error, they will be identified as different.

If at least one genome is heterozygous, (01x11 or 01x00 or 01x01 or 00x01 or 11x01), then the probability that the two pseudo haploid genomes do not differ is always equal to ½ because one allele will always be 0 (or 1) and the other will have a probability ½ to be 0 or 1.

The implementation in SPLATCHE thus consist in counting the number of differences between each pair of simulated diploid genomes by looping over all positions and 1) draw a random number RN between 0 and 1, 2) compare RN to the corresponding $P_{nodiff}$ from the above table depending on both genotypes; 3) if RN > $P_{nodiff}$ add one nucleotide difference between the two genomes. At the end of the loop the number of differences is divided by the total number of compared positions.

Note that this equation is not taking into account post-mortem damage (PMD), which would increase the error rate a little. However, as the goal is not to estimate this error rate but to take it into consideration in the analyses by using a prior distribution, it should not bias the results as the effect of PMD would be included in the prior range.

### Supplementary Data

**Supplementary Data 1. Information of the genomic data used in the present study.** The information provided are: the name of the genome (Genome); if the individual was classified as being a hunter-gatherer (HG) or a farmer (FA) (Population_Group); the latitude of the area where the individual was found (Latitude); the longitude of the area where the individual was found (Longitude); the age of the genome in years before present (Mean_Age_YBP); the population sample to which the genome was assigned for estimating the mean intra- and inter-sample pseudo-haploid nucleotide diversity and performing the simulations (Sample); the study which produced the genome (Publication). The full references of the publications from which the genomes were retrieved are also provided.

**Supplementary Data 2. Pairwise values of pseudo-haploid nucleotide diversity.** The values of pseudo-haploid nucleotide diversity for all genome pairs are provided.

**Supplementary Data 3. Mean values of pseudo-haploid nucleotide diversity.** The mean values of intra- and inter-sample pseudo-haploid nucleotide diversity are provided. For creating the samples, genomes sampled in the same deme and at the same generation during the simulations were grouped together. The genomes belonging to each sample can be found in Supplementary Data 1.
